# phyloDS: phylogeny-aware differential splicing across species, beyond reference annotations

**DOI:** 10.64898/2026.10.05.756841

**Authors:** Arnaud Liehrmann, Louis Carrel Billiard, Paul Bastide, Mélina Gallopin, Hugues Richard, Élodie Laine

## Abstract

Alternative splicing diverges rapidly between species, yet its comparative analysis lags far behind that of gene expression: junction orthology is defined on a single reference genome, species are treated as independent samples, and detected shifts are rarely grounded in sequence-level evidence. We present phyloDS, an RNA-seq–driven framework that resolves all three limitations. phyloDS defines orthologous splice junctions on multiple sequence alignments of one-to-one orthologous genes, tests their differential usage with a regularized Ornstein–Uhlenbeck model, and cross-validates detected shifts against splice-site predictions made from genomic sequence alone. Applied to primate brain and rodent heart transcriptomes, phyloDS recovered up to 2.9-fold more shared orthologous splice junctions than a reference-based strategy and kept type I error near nominal where competing methods inflate it more than two-fold. It identified 234 and 502 differentially used orthologous splice junctions in *hominoids* and *mole-rats* (86 and 141 unannotated), revealing a mitochondrial and contractile splicing signature in *mole-rats*.

## 1 Introduction

Alternative splicing is a pervasive feature of eukaryotic genomes. The vast majority of multi-exonic genes produce more than one transcript isoform [Pan et al., 2008, Wang et al., 2008, Nilsen and Graveley, 2010], and large-scale comparative studies have established that splicing patterns diverge rapidly along evolutionary lineages [Barbosa-Morais et al., 2012, Merkin et al., 2012, Reyes et al., 2013, Gueroussov et al., 2015]. By generating tissue-, stage-, and condition-specific isoforms from a shared genomic substrate, alternative splicing constitutes a major axis of post-transcriptional regulation and a plausible reservoir for phenotypic divergence between species [Xia et al., 2024, Wright et al., 2022, Verta and Jacobs, 2022, Hunter and Xing, 2026]. This makes it a natural target for comparative transcriptomics. Yet this potential has been far less exploited than differential gene expression in comparative analyses between species [Brawand et al., 2011, Perry et al., 2012, Berthelot et al., 2017, Stern and Crandall, 2018, Cardoso-Moreira et al., 2019, El Taher et al., 2020, Chalopin et al., 2026].

This asymmetry reflects three challenges that comparative gene expression has resolved through mature foundations, but that comparative splicing has yet to address. The first two stem from the same fact: species are connected by a shared evolutionary history that shapes both sequence and expression divergence. Sequence divergence makes identifying orthologous units across distantly related species non-trivial. This is a long-standing, well-studied problem at the gene level, for which robust solutions exist [Koonin, 2005, Emms and Kelly, 2019, Kirilenko et al., 2023]. Expression divergence means that observed variation may merely reflect shared phylogenetic history rather than *bona fide* lineage-associated shift, a problem now addressed by statistical frameworks that explicitly model phylogenetic dependence in expression data [Rohlfs and Nielsen, 2015, Gallopin et al., 2026]. The third challenge connects the first two: shifts detected at the expression or splicing level still need to be grounded in complementary sequence-level evidence, such as cis-regulatory landscape divergence, to enable cross-validation and mechanistic hypotheses [Daunesse et al., 2025].

Comparative splicing studies have sidestepped these challenges by reducing the multi-species problem to a single-species analysis at one or more steps, overlooking divergence at both levels. At the orthology step, sequence divergence is overlooked: splice junction orthology is defined in a reference-based manner, aligning RNA-seq reads from all species onto a single reference genome and testing alternative splicing events pre-annotated in that reference [Recinos et al., 2024]. As a result, splice junctions diverging beyond what the alignment recovers, or escaping the reference annotation, become invisible. At the testing step, usage divergence is overlooked: samples from all species are treated as independent [Xiong et al., 2018, Mazin et al., 2021, Ritter et al., 2024], even though shared ancestry correlates splice junction usage along the phylogeny, and pooling them as if independent inflates type I error rates, as previously shown for differential gene expression [Bastide et al., 2023]. Finally, none of the studies systematically integrates detected shifts with orthogonal sequence-level evidence for cross-validation and hypothesis generation. While this body of work has produced important biological insights, the field still lacks a solution that jointly addresses the three challenges.

We present phyloDS, an RNA-seq driven method for differential splicing analysis between two groups of species. Rather than reducing the multi-species problem to a single-species analysis, as done by most comparative splicing studies to date, phyloDS resolves splice junction orthology, accounts for phylogenetic non-independence in usage, and grounds detected shifts in sequence-level evidence – jointly. At the core of phyloDS is a simple statistical object: the within-sample log-ratio between a focal junction and its competing junctions, which decouples splicing from gene expression and is intrinsically normalized for library size. To define orthologous splice junctions (OSJs) in an unbiased manner, phyloDS calls splice junctions directly from species-specific RNA-seq data and establishes their orthology using a shared coordinate system built from genomic sequence alignments of orthologous genes, rather than anchoring them to a single reference genome. To model phylogenetic dependence among species, phyloDS builds on phyloDE [Gallopin et al., 2026], a regularized phylogenetic model for correlated count data. To support mechanistic hypotheses, phyloDS quantifies the agreement between transcriptomic and genomic divergence by comparing RNA-seq–inferred differential OSJ usage with splice site strength predictions obtained from genomic sequence alone using OpenSpliceAI [Chao et al., 2025], and exports alignment-anchored Integrative Genomics Viewer (IGV) sessions [Robinson et al., 2011] that integrate statistical, sequence, and annotation evidence within a shared cross-species coordinate system.

We applied phyloDS to two phylogenetically structured RNA-seq datasets that pair phylogenetic structure with striking, well-studied phenotypic contrasts, spanning different levels of evolutionary divergence. In primates, the hominoid lineage shows a marked expansion in brain size and cognitive function relative to other primates [Smaers and Vanier, 2019], a contrast that has driven substantial interest in the genomic and transcriptomic basis of human and great-ape brain evolution. A previous study of brain transcriptomes from eight primate species identified over a thousand exons with lineage-specific splicing shifts genome-wide, but relied on a reference-based approach, aligning all species to the human genome [Recinos et al., 2024]. Applying phyloDS to the same transcriptomes, augmented with one additional great-ape species [Xu et al., 2018], we identified 234 differentially used OSJs across 158 genes, including 86 unannotated OSJs inaccessible to reference-based detection, and recovered the well-characterized *MAPT* exon 10/exon 2 switch. In rodents, the mole-rat lineage is an emerging model system prized for its extreme tolerance to hypoxia [Ivy et al., 2020], hypercapnia [Park and Reznick, 2023], and oxidative stress [Lewis et al., 2013], and has motivated recent studies of gene expression and regulatory evolution [Daunesse et al., 2025]. To our knowledge, however, alternative splicing has never been compared between mole-rats and other rodents. Applied to four-rodent heart transcriptomes, phyloDS identified 502 differentially used OSJs across 322 genes, including 141 unannotated, strongly enriched for mitochondrial and contractile programs – a splicing signature that aligns closely with the mole-rats’ known physiology. This analysis thus constitutes the first evidence that alternative splicing contributes to mole-rats’ physiological adaptations. In both datasets, differentially used OSJs showed strong directional concordance with genomic sequence-only predictions from OpenSpliceAI, supporting a cis-regulatory basis for these changes.

On all three challenges, phyloDS delivers where existing strategies fall short. First, its alignment-based orthology recovered 2.9 times more OSJs shared across all species than a reference-based strategy in the more divergent *mole-rats* dataset, capturing annotated orthologous splicing events while uncovering hundreds of novel ones. Second, its regularized phylogenetic model, implemented in phyloDE, keeps type I error rates close to the nominal level where competing methods inflate more than two-fold. Third, its cross-species coordinate system grounds detected differentially used OSJs in sequence-level evidence and turns genome-wide lists into cis-regulatory hypotheses.

phyloDS provides a robust, versatile, and practical framework to detect, validate, visualize and interpret evolutionary changes in alternative splicing from RNA-seq data. It is implemented as an R package available at https://github.com/aLiehrmann/phyloDS.

## 2 Results

### 2.1 phyloDS detects differentially used OSJs and supports their interpretation in three stages: summarize, test, explore

phyloDS enables differential analysis of OSJ usage between two predefined groups of species by integrating RNA-seq data, sequence conservation, and phylogenetic information. The method takes as input species-specific genome sequences, an ultrametric phylogenetic tree, group membership information, genomic coordinates of one-to-one orthologous genes, and processed RNA-seq alignments (BAM files). The method then unfolds in three stages, illustrated in Figure 1.

**Figure 1:**
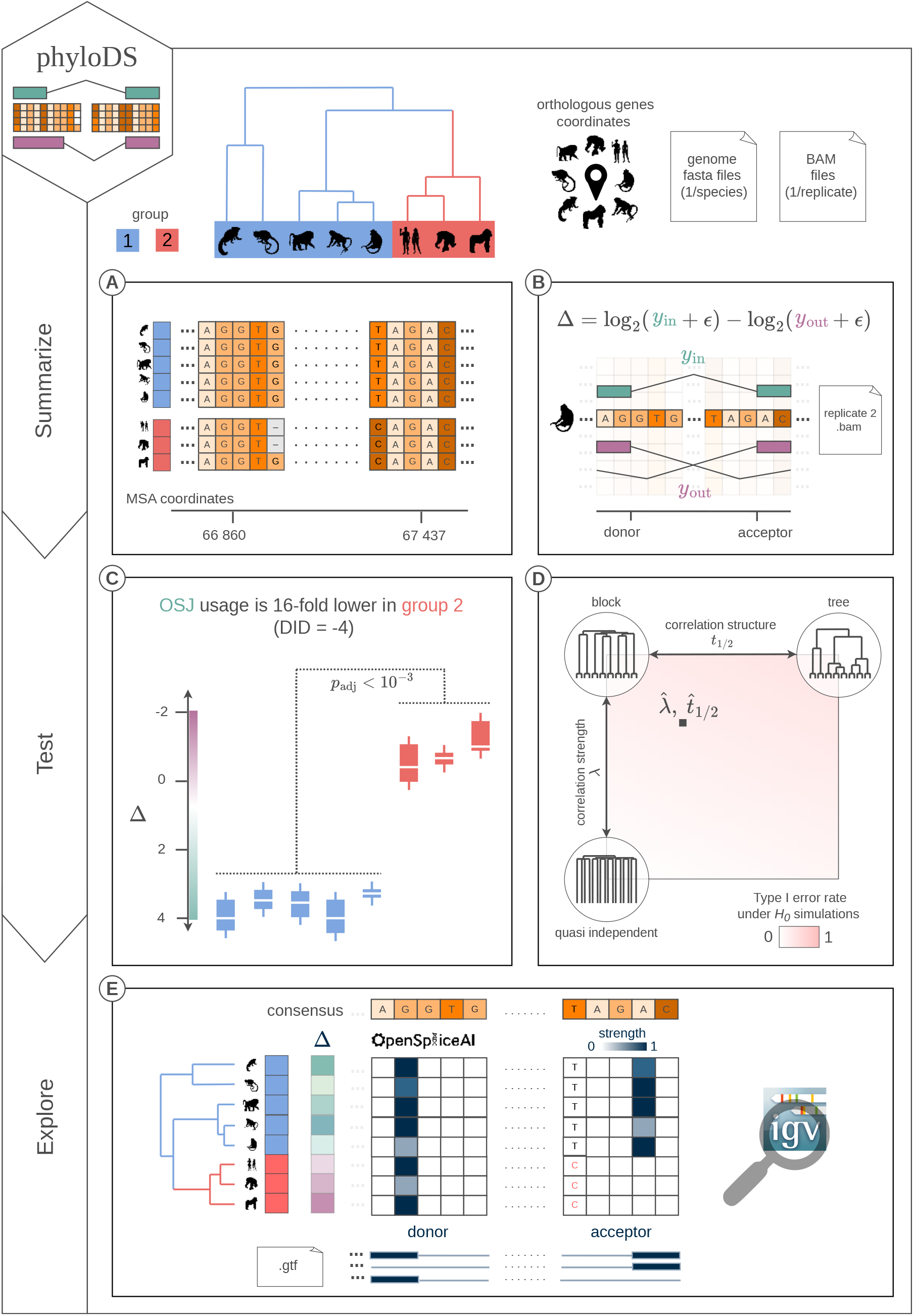
Overview of phyloDS. Inputs. (top): an ultrametric species tree whose tips are assigned to two groups (group 1 in blue, group 2 in red; here five non-hominoid primates versus three hominoids), coordinates of 1-to-1 orthologous genes, one genome FASTA file per species, and one RNA-seq BAM file per replicate. The same colour code for groups is used throughout the figure. **Summarize** (**A**, **B**). (**A**) Orthologous splice junctions (OSJs) are defined in the coordinate system of a multiple sequence alignment of the orthologous genes. A dash marks a gap, that is, a site absent from a species. A donor and an acceptor alignment column together define one OSJ. (**B**) For every replicate, usage of a focal OSJ is quantified from split reads in the corresponding BAM file as the within-sample log-ratio Δ between the focal junction (*y*_in_, teal) and the aggregate of its competing junctions (*y*_out_, purple), with pseudocount *ɛ*. **Test** (**C**, **D**). (**C**) Per-species distributions of Δ for one OSJ. The group contrast, estimated with phyloDE while accounting for phylogenetic covariance, is the difference-in-differences (DID), reported with a Benjamini–Hochberg adjusted *p*-value. (**D**) Calibration. The two Ornstein–Uhlenbeck parameters span a plane: *λ* (vertical) sets the strength of the correlation, from quasi-independent replicates at the bottom, and *t*_1*/*2_ (horizontal) sets its structure, from species blocks on the left to the full tree topology on the right; the circular insets show the correspondingly transformed trees. The background shows the type I error rate measured on *H*_0_ simulations, and the black square marks the parameters estimated on the dataset. **Explore** (**E**, **F**). (**E**) Each differentially used OSJ is placed in its sequence context: OpenSpliceAI donor and acceptor strengths predicted per species from genomic sequence alone (blue scale), the aligned nucleotides flanking both splice sites, and exon annotations projected from the species GTF files. (**F**) All of this is exported as a ready-to-use IGV session whose tracks bundle the outputs of steps **A**, **C** and **E** within the shared cross-species coordinate system.

#### Summarize: phyloDS maps species-specific splice junctions onto a shared cross-species coordinate system to define OSJs

phyloDS first extracts genomic sequences corresponding to orthologous genes and aligns them to define a single coordinate system common to all species (Figure 1.A). It then identifies splice-junction donor and acceptor sites from BAM files and maps them onto the multiple sequence alignment (MSA). This step defines OSJs and quantifies spliced-in and spliced-out read counts in a consistent cross-species framework (Figure 1.B).

#### Test: phyloDS tests OSJs for differential usage while controlling phylogenetic false positives

phyloDS tests each OSJ for differential usage between the two species groups, delegating this step to phyloDE. phyloDE estimates and regularizes a phylogenetically informed covariance structure, modeled as an Ornstein-Uhlenbeck (OU) process, decorrelates the data, then applies empirical Bayes shrinkage to compute one moderated *t*-statistic and corresponding *p*-value per OSJ (Figure 1.C). To verify that this controls phylogenetic false positives, phyloDS can also simulate realistic OSJ counts, capturing both RNA-seq overdispersion and evolutionary dependence, and assess its type I error rate as two correlation parameters increase: species-structured correlation among replicates and tree-structured correlation among species (Figure 1.D).

#### Explore: phyloDS annotates, visualizes, and links differentially used OSJs to sequence-level evidence

Finally, phyloDS annotates differentially used OSJs from the exon boundaries of one or more user-designated species, and flags those that involve unannotated splice sites. In addition, it provides species-specific predictions of splice-site strength, computed using OpenSpliceAI, to help users assess whether differentially used OSJs are supported by underlying genomic sequence variation nearby splice donor and acceptor sites. All results are then exported in visualization-ready formats compatible with IGV, enabling joint inspection of OSJs, annotations, and splice-site strengths along the MSA (Figure 1.E).

A detailed description of the phyloDS workflow is provided in the Methods section (see *phyloDS workflow and implementation*).

### 2.2 Applied to the *hominoids* and *mole-rat* datasets, phyloDS enables genome-wide detection of differentially used OSJs and provides supporting sequence-level evidence

We applied phyloDS to two phylogenetically structured RNA-seq datasets spanning different levels of evolutionary divergence. The first dataset (*hominoids*) comprises brain samples from eight primate species diverged over approximately *∼* 43 million years [Kumar et al., 2017], each represented by at least two biological replicates (Figure 2.A, left). The second dataset (*mole-rats*) includes heart samples from four rodent species spanning a substantially deeper divergence of *∼* 70 million years [Kumar et al., 2017], each represented by at least five biological replicates (Figure 2.A, right). In both datasets, the contrasts of interest define two groups aligned with the phylogeny: group 2 corresponds to the focal lineage (three hominoids or two mole-rats), whereas group 1 comprises the remaining species used as outgroup. Quality-control analyses, including principal component analysis and histograms of raw *p*-values, did not reveal any anomalies in either dataset (Figure S1).

**Figure 2:**
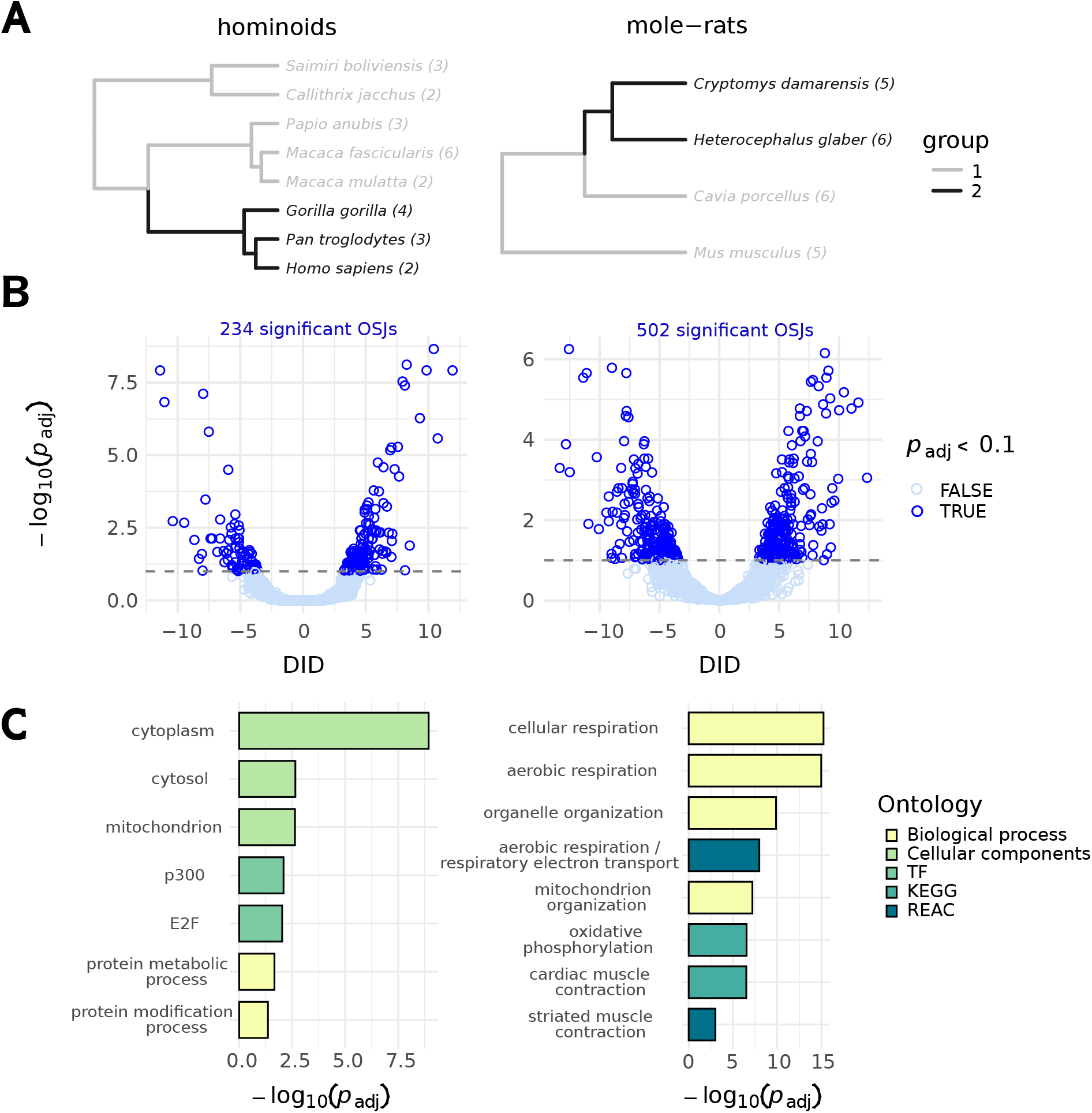
Application of phyloDS to two phylogenetically structured RNA-seq datasets. (**left**) *hominoids* brain dataset. (**right**) *mole-rats* heart dataset. (**A**) Species phylogeny and study design: group 1 (grey branches) and group 2 (dark branches); numbers in parentheses indicate the number of biological replicates per species. (**B**) Volcano plot summarizing differential OSJ usage; 234 OSJs were identified as differentially used (*p*_adj_ *<* 0.1) in *hominoids* and 502 in *mole-rats*. The *x*-axis shows the phylogenetically corrected difference in OSJ usage on the log_2_ scale (DID), estimated with phyloDE; positive values indicate higher relative usage in group 2 than in group 1. (**C**) Functional enrichment analysis of genes harboring at least one significant OSJ. In the *hominoids* brain dataset, enriched terms correspond to broadly defined intracellular and protein regulatory processes. In the *mole-rats* heart dataset, enrichment highlights pathways related to mitochondrial bioenergetics and cardiac contractile machinery.

#### phyloDS reveals hundreds of differentially used OSJs in *hominoids* and *mole-rats*

Overall, phyloDS identified 234 differentially used OSJs (Benjamini–Hochberg-adjusted *p*-values *<* 0.1) across 158 genes in *hominoids* (Figure 2.B, left; Table S1) and 502 across 322 genes in *mole-rats* (Figure 2.B, right; Table S2). Effect sizes (absolute DID – the phylogenetically corrected log_2_ difference in OSJ usage between groups) ranged from 3.288 to 11.914 (median 4.861) in *hominoids*, and from 3.324 to 13.336 (median 5.281) in *mole-rats*.

#### A substantial fraction of differentially used OSJs involve previously unannotated splice sites

phyloDS recovered 86 (*hominoids*; Table S3) and 141 (*mole-rats*; Table S4) differentially used OSJs involving at least one unannotated splice site (either 5’ or 3’) in species from the group with increased OSJ usage. In *hominoids*, illustrative examples include a skipping exon (SE) in *USP31*, an alternative 5’ splice site (A5) in *UBE4A*, an alternative last exon (AL) in *RB1CC1*, and an alternative 3’ splice site (A3) in *NCKIPSD* (Figures S2–S5). In *mole-rats*, additional examples include an alternative first exon (AF) in *PHLDB1* and an SE in *IMMT* (Figures S6–S7). These results show that combining MSA-based orthology with RNA-seq evidence enables the detection of differentially used OSJs, even when splice sites are missing from current annotations.

#### Differentially used OSJs map broadly onto protein regulation in *hominoids*, and sharply onto mitochondrial and contractile programs in *mole-rats*

To further assess whether differential OSJ usage is associated with coherent biological programs, we performed functional enrichment analyses on genes with at least one differentially used OSJ. In the *hominoids* brain dataset, enrichments were detectable but general at the GO level (Figure 2.C, left; Table S5), with over-representation of broad categories related to *protein metabolic/modification processes* (GO:0019538, GO:0036211; *p*_adj_ = 6 *×* 10*^−^*^3^) and subcellular localization biased toward *cytoplasm/cytosol/mitochondrion* (GO:0005737, GO:0005829, GO:0005739; *p*_adj_ *≤* 6.7 *×* 10*^−^*^3^). Transcription factor motif enrichments were dominated by E2F-family and co-activator motifs (e.g., *E2F/E2F-2/E2F-4* and *p300*; *p*_adj_ = 7 *×* 10*^−^*^3^), consistent with a bias toward genes involved in macromolecule and protein regulation.

In the *mole-rats* heart dataset (Figure 2.C, right; Table S6), enrichments were strongly dominated by mitochondrial and energetic functions, including *organelle/mitochondrion organization* (GO:0006996 / GO:0007005; *p*_adj_ *≤* 3 *×* 10*^−^*^10^) and *cellular/aerobic respiration* (GO:0045333 / GO:0009060; *p*_adj_ = 4.07 *×* 10*^−^*^13^). Consistently, pathway databases highlighted *oxidative phosphorylation* (KEGG:00190; *p*_adj_ = 5.55 *×* 10*^−^*^7^) and *cardiac muscle contraction* (KEGG:04260; *p*_adj_ = 1.91 *×* 10*^−^*^6^). Reactome provided a concordant signal for *aerobic respiration / respiratory electron transport* (e.g., REAC:R-MMU-1428517; *p*_adj_ = 2.4 *×* 10*^−^*^8^) and *striated muscle contraction* (e.g., REAC:R-MMU-390522; *p*_adj_ *≤* 1.38 *×* 10*^−^*^3^). These enrichments suggest that mole-rat-associated splicing shifts in heart preferentially affect genes central to mitochondrial bioenergetics and contractile physiology.

#### Differentially used OSJs are strongly concordant with sequence-level evidence, supporting a cis-regulatory basis

We further evaluated whether differentially used OSJs were supported by underlying genomic sequence variation. For a set of representative OSJs, donor and acceptor splice site strengths were predicted independently in each species with OpenSpliceAI and summarized through a sequence-based 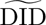 estimate (see the *Comparison with OpenSpliceAI–inferred* 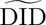 section in Methods). If the observed shifts in OSJ usage reflect cis-regulatory changes (mutations, indels) affecting splice site recognition, the direction of the RNA-seq and sequence-based effects should be concordant. Consistent with this expectation, differentially used OSJs detected by phyloDS (*p*_adj_ *<* 0.1) showed a strong enrichment for concordant effect directions with the sequence-based 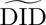 estimates (Figure 3.A). In the *hominoids* dataset, Fisher’s exact test yielded an odds ratio of 17.92 (*p*_adj_ = 9.32 *×* 10*^−^*^9^). In the *mole-rats* dataset, concordance was even stronger, with an odds ratio of 29.02 (*p*_adj_ = 2.99 *×* 10*^−^*^28^). No such association was found for clearly non-differentially used OSJs (*p*_adj_ *>* 0.9; Figure S11), confirming that the concordance is specific to phyloDS-detected differentially used OSJs and consistent with a cis-regulatory basis.

**Figure 3:**
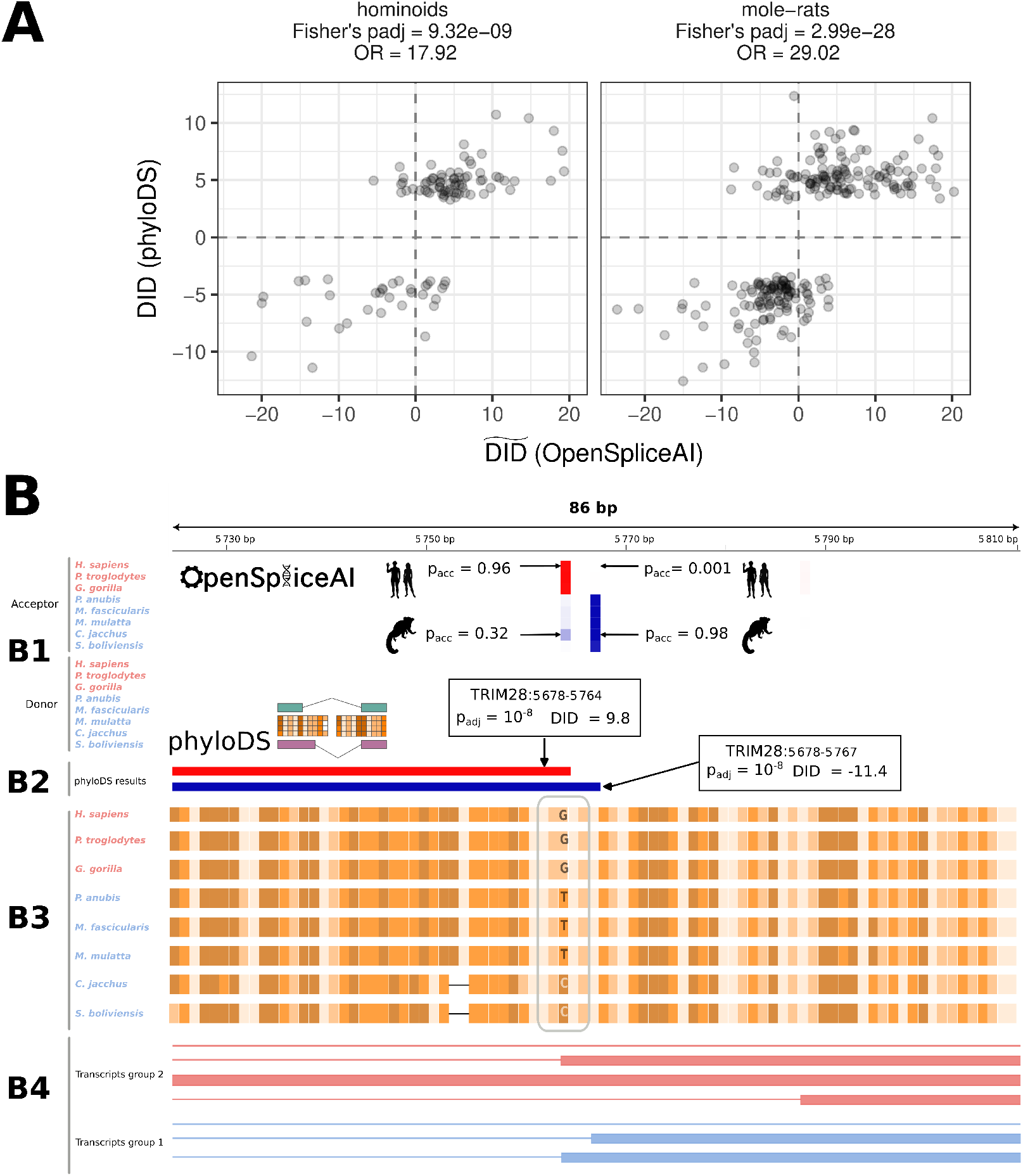
(**A**) Comparison of difference-in-differences estimates derived from RNA-Seq data using phyloDS and from sequence-based predictions using OpenSpliceAI. Panels are stratified by dataset (*hominoids* and *mole-rats*). Each point represents one OSJ, as identified by phyloDS (*p*_adj_ *<* 0.1). Concordance between RNA-Seq and sequence-based effect directions is highly significant in both datasets according to Fisher’s exact test. The corresponding analysis for clearly non-differentially used OSJs (*p*_adj_ *>* 0.9) is presented in Figure S11, where Fisher’s exact test is not significant. (**B**) IGV session exported by phyloDS illustrating differential usage of an A3 event in *TRIM28* (exon 12; ENSE00003513051) between hominoids and other primates. In *Homo sapiens*, the canonical acceptor is located at position 58, 548, 862, with an alternative acceptor at +3. (**B1**) OpenSpliceAI splice site strength tracks (acceptors shown above; donors below). Darker colors (red for group 2, hominoids; blue for group 1, other primates) indicate higher predicted splice site strengths, whereas white correspond to values close to or equal to zero. (**B2**) Differentially used OSJ (*p*_adj_ *<* 0.1) are shown as colored rectangles: red indicates higher relative usage in group 2 compared to group 1 (DID *>* 0), and blue indicates the opposite (DID *<* 0). Grey spans (not shown here) correspond to non-differentially used OSJs (*p*_adj_ *>* 0.1). (**B3**) MSA providing a shared coordinate system across species; the boxed region highlights the local AG dinucleotide context surrounding the competing acceptor sites, where the central G observed in hominoids is replaced by T or C in other primates. (**B4**) Group-specific transcript annotations mapped onto the MSA coordinates. Exons are shown as thick lines and introns as thin lines. The left boundary of each exon corresponds to the 3’ splice site (acceptor), and the right boundary to the 5’ splice site (donor; not shown here).

### 2.3 phyloDS MSA-anchored IGV sessions place differentially used OSJs in their cross-species sequence context

Beyond producing genome-wide lists of differentially used OSJs, phyloDS exports a ready-to-use IGV session that integrates statistical evidence, sequence context, and species annotations within a shared evolutionary coordinate system defined by the MSA. Figure 3.B illustrates this session for an A3 event in *TRIM28* showing differential usage in the *hominoids* dataset. Two differentially used OSJs share the same donor (not shown on the figure) but use acceptors separated by 3 nucleotides (nt). In the OSJ-span track (Figure 3.B2), the upstream acceptor shows DID = 9.8 (increased usage of the OSJ in hominoids compared to other primates), whereas the downstream acceptor shows DID = *−*11.4 (decreased usage in hominoids), consistent with a clade-specific switch. Sequence-based predictions from OpenSpliceAI (Figure 3.B1), which provide an orthogonal validation, mirror this pattern. In hominoids, the upstream acceptor site exhibits a high predicted strength, whereas the downstream site is predicted to be weak (for example, 0.96 and 0.001, respectively, in *Homo sapiens*). In contrast, outgroup species display the opposite configuration (for example, 0.32 and 0.98, respectively, in *Callithrix jacchus*). The MSA track (Figure 3.B3) anchors both acceptors in a shared evolutionary frame and highlights the local AG context (dark rectangle). The 3-nt displacement corresponds to a one-codon shift, generating a single glycine residue difference between proteoforms. Consistent with the mechanistic model of Jayasinghe et al. [2018], mutation of the central nucleotide within an agGag-like context can bias competition between adjacent AG sites, providing a plausible sequence-level explanation for the observed evolutionary switch. Notably, in this example, projected transcript annotations (Figure 3.B4) are also coherent with this pattern: hominoids (*Homo sapiens*, *Pan troglodytes*, *Gorilla gorilla*) contain only the upstream acceptor annotation, whereas other primates retain annotations for both upstream and downstream sites. Because phyloDS defines OSJs without reference to any annotation, this agreement provides a further, independent validation of the detected switch. This example illustrates how phyloDS enables a direct transition from a significant DID statistic to a testable, sequence-supported hypothesis about the cis-regulatory basis of the switch.

### 2.4 phyloDS enables an unbiased definition of OSJs between species and controls for phylogeny-related false positives

To our knowledge, no comparative splicing study to date provides a dedicated package for testing differential splicing between groups of species, so a head-to-head comparison of software is not possible. We therefore decomposed published pipelines into the three design choices that separate them from phyloDS (how splice junction orthology is established, at which granularity junctions are summarized before testing, and how the test accounts for phylogenetic dependence) and benchmarked each choice in turn (Figure 4.A, Figure 4.B and Figure 5, respectively). The closest comparator is Recinos et al. [2024], which addresses the same question and makes the alternative choice on all three axes: RNA-seq reads from every species are aligned to a single reference genome, usage is summarized at the level of SE events annotated in that reference, and lineage shifts are tested with eve [Rohlfs and Nielsen, 2015]. Two further studies [Mazin et al., 2021, Ritter et al., 2024] treat samples from different species as independent replicates, although neither tests two groups of species as such. We therefore included limma, a non-phylogenetic linear model, in our comparison (Figure 5).

**Figure 4:**
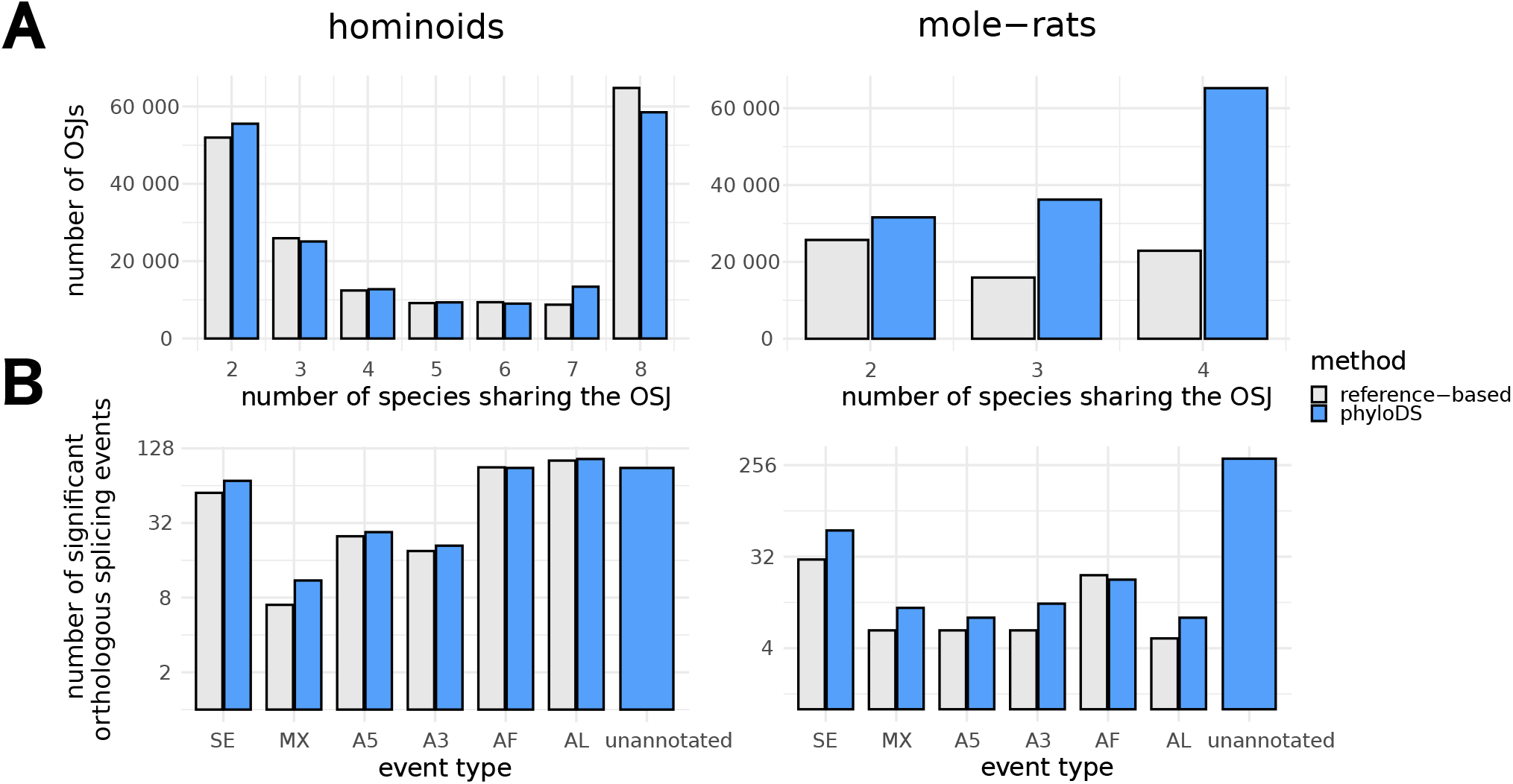
Comparison between phyloDS and reference-based strategy. (**A**) Number of OSJs inferred across species using either the MSA-based strategy implemented in phyloDS or a reference-based strategy in which RNA-seq reads from all species were aligned to a single reference genome (*Homo sapiens* for *hominoids*, *Mus musculus* for *mole-rats*). In the shallowly diverged *hominoids* dataset, both strategies recover comparable numbers of OSJs, whereas in the more divergent *mole-rats* dataset, phyloDS substantially improves recovery of OSJs. (**B**) Number of differentially used orthologous splicing events detected using either the annotation-free OSJ-level testing strategy implemented in phyloDS or a reference-based event-level testing strategy, i.e., using annotated splicing events from the reference species. On the annotated events, both strategies detect comparable numbers of differentially used orthologous events, whereas phyloDS additionally detects hundreds of differentially used unannotated orthologous events.

**Figure 5:**
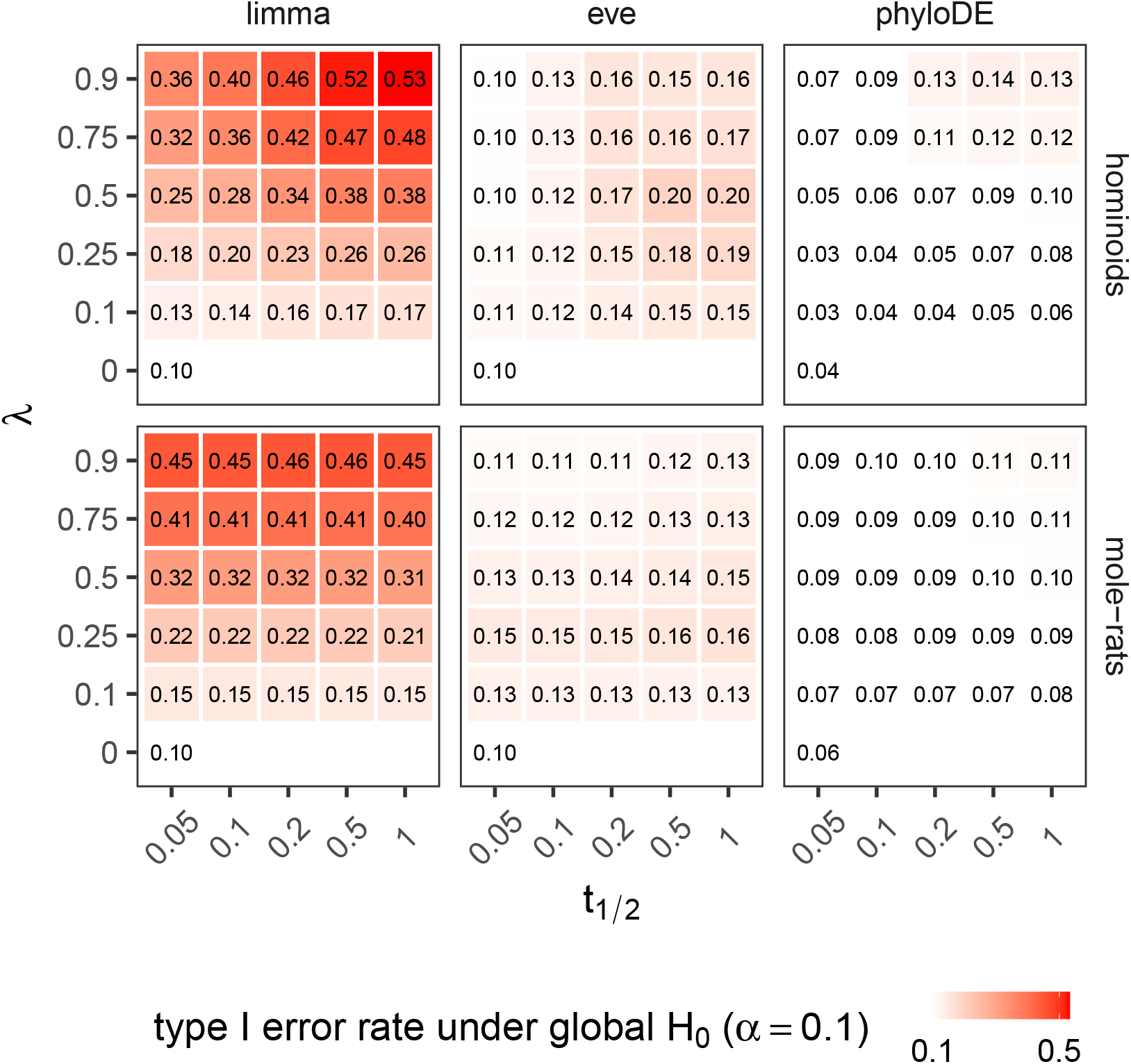
Type I error rate under the global null hypothesis (*H*_0_) at nominal level *α* = 0.1. OSJ counts (spliced-in and spliced-out) were simulated with compcodeR under a phylogenetic Poisson log-normal model calibrated on the *hominoids* and *mole-rats* datasets to match empirical mean–dispersion relationships. For each scenario, 5 000 OSJs were simulated with no group effect (DID = 0), across a grid of phylogenetic signals (*λ*; ranging from independent replicates to species-structured correlations) and OU half-lives (*t*_1*/*2_; ranging from independent species to tree-structured correlations), with 50 replicates. The empirical type I error rate was computed as the proportion of raw *p*-values below *α* and averaged across replicates. Methods shown include a regularized non-phylogenetic linear model approach (limma), a phylogenetic approach that estimates parameters independently for each OSJ (eve), and phyloDE, which forms the core of phyloDS and combines phylogenetic modeling with cross-OSJ regularization. phyloDE is the only approach that consistently maintains type I error rates close to the nominal level across scenarios and datasets. Results for two additional methods, limmaCorr and phylolm, are shown in Figure S10

#### phyloDS recovers more OSJs than reference-based strategy at deeper divergence

We compared two strategies to infer OSJs: the MSA-based strategy implemented in phyloDS and a reference-based strategy, in which OSJs are identified across species by aligning RNA-seq reads to a common reference genome (*Homo sapiens* for *hominoids* and *Mus musculus* for *mole-rats*; see *Comparison with reference-based OSJ inference* in Methods). In the *hominoids* dataset, which spans a relatively shallow phylogenetic divergence, the two strategies were comparable overall, with the reference-based strategy recovering slightly more all-species OSJs. The MSA-based strategy identified 183 651 OSJs supported in at least two species (versus 182 418 for the reference-based method) and 58 506 shared across all eight species (versus 64 797; Figure 4.A, left). In contrast, in the more evolutionarily divergent *mole-rats* dataset, the MSA-based strategy substantially improved OSJ recovery. It identified 133 014 OSJs supported in at least two species, compared with 64 547 using the reference-based method, and detected 65 212 OSJs shared across all four species versus 22 903, corresponding to a 2.9-fold increase (Figure 4.A, right). This result shows that our strategy of defining a common coordinate system achieves higher sensitivity for more divergent species than strategies inferring orthology using a single reference.

#### phyloDS recovers hundreds of differentially used unannotated orthologous splicing events without losing sensitivity on annotated ones

OSJ-level testing should capture splicing shifts that reference-based event-level testing misses, but testing each OSJ in isolation may sacrifice statistical power, a gain-versus-loss trade-off we set out to quantify. Two axes distinguish these strategies: the unit being tested (a single OSJ vs. a group of OSJs aggregated into an event) and the reference frame used to define it (annotation-free, based on multi-species alignment, vs. reference-based, anchored on a single species’ annotation). phyloDS implements an annotation-free, OSJ-level strategy, which we compared to a reference-based, event-level strategy that jointly tests groups of OSJs corresponding to annotated splicing events from a reference species (*Homo sapiens* in *hominoids* and *Mus musculus* in *mole-rats*). For example, a SE event aggregates three OSJs: two corresponding to inclusion of the exon and one to its exclusion (see *Comparison with reference-based differential splicing event detection* in Methods). In the *hominoids* dataset, phyloDS OSJ-level testing recovered numbers of differentially used events of all types comparable to those recovered by reference-based event-level testing: AF and AL were the most abundant, followed by SE, A3, A5, and finally MX (mutually exclusive exons; Figure 4.B, left). A similar pattern was observed in the *mole-rats* dataset (Figure 4.B, right). In addition, phyloDS detected differentially used unannotated events across species that reference-based event-level testing cannot access: 89 in *hominoids* and 296 in *mole-rats*. These results show that OSJ-level testing in phyloDS, which is unbiased with respect to reference-species annotations, matches the sensitivity of reference-based event-level testing while additionally detecting hundreds of differentially used unannotated orthologous events.

#### phyloDS controls type I error rates where competing methods inflate more than two-fold

To test whether our phyloDE-based strategy keeps the type I error rate at its nominal level when there is no true group effect (DID = 0), we ran empirical null simulations under a phylogenetic Poisson log-normal model calibrated separately on each dataset (see *Empirical null simulations with compcodeR* in Methods). We simulated OSJ profiles with DID = 0 across a grid of two correlation parameters: phylogenetic signal *λ*, from independent replicates (*λ* = 0) to strongly correlated replicates (*λ* = 0.9), and OU half-life *t*_1*/*2_, from block-structured correlations (*t*_1*/*2_ = 0.05) to tree-structured correlations (*t*_1*/*2_ = 1; Figures S8–S9). We evaluated alternative inference strategies, including limma, limmaCorr, phylolm, and eve (see *Model inference* in Methods). In contrast to phyloDE, both limma and limmaCorr ignore phylogenetic correlations, while phylolm and eve estimate phylogenetic parameters independently for each OSJ without regularization. Type I error was quantified as the proportion of raw *p*-values below the nominal threshold *α* = 0.1, averaged over 50 replicates per scenario.

phyloDE was the only method to keep the type I error rate near the nominal *α* = 0.1 across the grid, while competing methods inflated in the evolutionary regimes estimated from the real data. For *hominoids*, around phyloDE’s global, robust trimmed-mean estimates (*λ̂*= 0.49, *t̂*_1*/*2_ = 0.31; see *Model inference*), eve reached type I error rates of 0.17–0.20 and the non-phylogenetic methods inflated even more (*>* 0.20), whereas phyloDE stayed calibrated (0.07–0.09; Figure 5, top panel). Consistently, for *mole-rats*, around the same estimates (*λ̂* = 0.74, *t̂*_1/2_ = 0.08), eve showed mild inflation (0.12) and the other methods’ inflation was substantial (*>* 0.16), whereas phyloDE remained below nominal (0.09; Figure 5, bottom panel). Together, these results show that combining OU-based phylogenetic modeling with cross-OSJ regularization, as in phyloDE, yields well-calibrated inference under realistic evolutionary regimes.

## 3 Discussion

Comparative transcriptomics routinely asks which genes changed expression along a lineage; it has had no equivalent way to ask which splice junctions changed usage. phyloDS closes that gap. To our knowledge, it is the first framework to address splice junction orthology, phylogeny-aware testing and sequence-level support within a single cross-species coordinate system, and to do so on real datasets spanning contrasting levels of evolutionary divergence. Each of them, as we discuss below, still leaves clear room for improvement.

phyloDS models a single statistical object: the within-sample log-ratio between a focal OSJ and its competing OSJs (Δ). Δ decouples splicing from gene expression, since global changes in transcript abundance leave it unchanged, and is intrinsically normalized for library size, being computed within each sample (see *Phylogenetic model for the differential usage of splice junctions* in Methods).

Applied to primate brain transcriptomes, phyloDS recovered differentially used OSJs in genes enriched for protein regulatory processes, but also encompassed established cases. Two differentially used OSJs in microtubule-associated protein tau (*MAPT*) map precisely to exon 2 and exon 10 (Table S1), whose anticorrelated inclusion underwent stepwise, lineage-specific changes along the catarrhine lineage, culminating in hominoids and reshaping tau isoform composition [Recinos et al., 2024]. In rodent heart transcriptomes, the signal was more focused on mitochondrial bioenergetics and cardiac contraction. One example is the cardiac troponin I gene *TNNI3*, whose splicing has been repeatedly modified in high-heart-rate mammals [Joyce et al., 2024], in which a differentially used OSJ indicates less frequent skipping of exon 4 in mole-rats (Table S2).

Working in the coordinate system of an MSA, rather than relative to a single reference genome, enables phyloDS to recover OSJs without relying on annotations, including at unannotated splice sites – and the gap over reference-based recovery widens as evolutionary divergence increases. This gain in recovery, however, may narrow at deep divergence, as nucleotide homology signals saturate at synonymous and intronic sites and become ambiguous in repetitive or indel-rich regions. Working at the nucleotide level also carries a computational cost: building the MSA is currently the main runtime bottleneck (*>* 97%) due to its quadratic complexity, requiring a length cap (150 kb) that excludes long loci. Evolutionary splicing graphs (ESGs) [Zea et al., 2021] offer a promising path forward. By aligning short orthologous exonic units across many genes and species, ESGs exploit amino-acid conservation beyond nucleotide signal saturation, and cluster sequences before alignment rather than constructing a single monolithic alignment for each locus – sidestepping both the saturation and the runtime bottleneck at once. Reconciling such protein-level orthology with the annotation-free, junction-level detection central to phyloDS is therefore a natural next step toward a more scalable and robust definition of orthologous splicing units.

On the statistical side, the regularized phylogenetic OU linear model of phyloDS keeps type I error close to nominal even with the handful of species and replicates per species typically available in comparative transcriptomics. With so few measurements, degrees of freedom are limited, making unregularized per-OSJ estimators such as phylolm or eve, as well as non-phylogenetic tools such as limma, prone to inflating type I error rates. This control, however, rests on one univariate quantity per OSJ, the change in the Δ (DID), testing each OSJ independently of the others. Yet a preliminary, test-free look at our own data suggests that clades may differ not only in mean usage but also in the way the focal and competing OSJs covary (Figure 6). Such patterns, together with the positive correlation expected among OSJs that share a *cis*/*trans* context or belong to the same transcript, call for joint rather than per-junction modelling. The phylogenetic empirical-Bayes framework recently proposed by Montoya et al. [2026] is a promising direction here. Such a model could allow phyloDS to test not only shifts in the mean usage of an OSJ but also changes in its covariance with other junctions, while scaling to the developmental and multi-tissue panels now available [Cardoso-Moreira et al., 2019, Mazin et al., 2021]. Modeling correlations among junctions may also help determine whether interspecific splicing variation reflects adaptation or is largely consistent with genetic drift and relaxed selection associated with effective population size [Bénitière et al., 2024].

**Figure 6:**
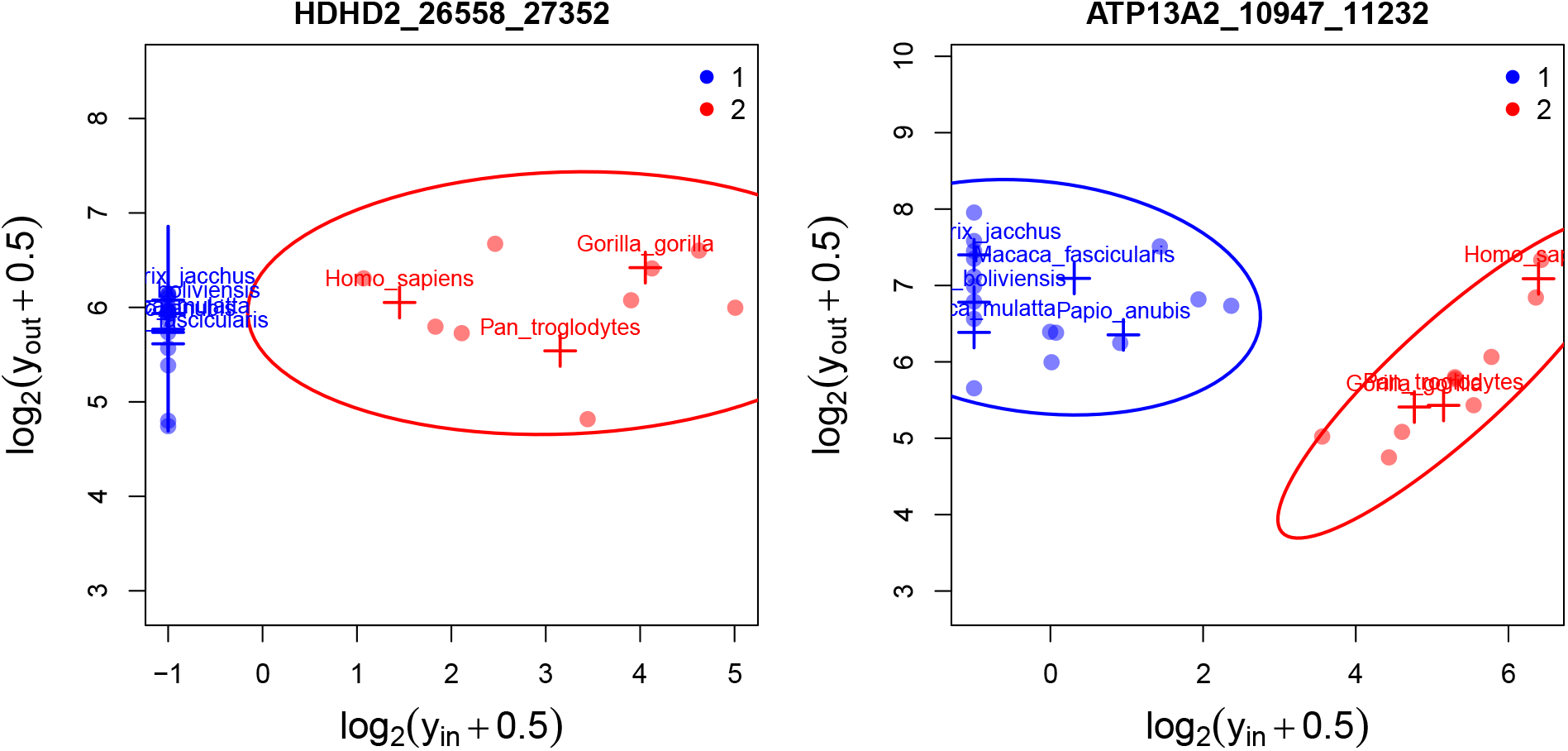
Clades may differ not only in relative usage but also in how the focal OSJ and its competitors covary. Two OSJs called differentially used by phyloDS in the *hominoids* dataset. Each point is a biological replicate, placed by its focal (*y*_in_) and competing (*y*_out_) log-counts (pseudocount 0.5); crosses mark species means, and ellipses summarize the within-clade covariance of hominoids (clade 2, red) and other primates (clade 1, blue). (**left**) In *HDHD2*, the focal OSJ is not detected in any sample from the other primates (*y*_in_ = 0, hence the vertical alignment at log_2_ 0.5 = *−*1), whereas in hominoids focal and competing log-counts show no clear covariation. (**right**) In *ATP13A2*, hominoids align along a narrow line roughly parallel to the diagonal, i.e. of near-constant Δ, as if their relative usage had become constrained, while the other primates remain decoupled. In both cases, phyloDS detects the shift in mean relative usage; the difference in covariance structure between clades is not modelled by the current per-OSJ test.

A last line of support comes from sequence itself. Across both datasets, the directional concordance between phyloDS effect sizes and OpenSpliceAI splice-site predictions, as exemplified by *TRIM28*, provides orthogonal sequence-level support for a cis-regulatory basis of many of these shifts, consistent with reports that interspecies splicing divergence is predominantly cis-directed [Barbosa-Morais et al., 2012]. This support is local by construction. OpenSpliceAI scores each splice site from a fixed 10,000-nt window of surrounding sequence. It therefore captures proximal cis signals but remains largely blind to distal exonic and intronic enhancers or silencers, as well as to trans-acting differences [Fu and Ares, 2014, Rogalska et al., 2024]. It is also trained on a single reference genome, such that predicted splice-site strengths and the effect-size scale of our sequence-based 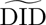 may be imperfectly calibrated across model and non-model species, particularly with increasing evolutionary divergence. Both limitations point to concrete next steps. Genome-native models such as AlphaGenome [Avsec et al., 2026] predict splicing from megabase-scale sequence context and may therefore capture more distal regulatory effects. In parallel, OpenSpliceAI was designed for transfer learning [Chao et al., 2025], and species-specific training should improve calibration with increasing distance from human or mouse and may ultimately allow Δ to be inferred from genome sequence alone, extending phyloDS to lineages lacking RNA-seq data.

phyloDS provides a robust, versatile, and practical framework for comparative transcriptomics, enabling the detection, validation, visualization, and interpretation of evolutionary changes in alternative splicing directly from RNA-seq data. By jointly addressing the challenges of orthology inference, phylogeny-aware statistical testing, and sequence-level validation, it transforms species-specific splice-junction observations into orthologous, phylogenetically corrected, and sequence-grounded units of analysis. In doing so, it places comparative splicing analysis on the same statistical footing as comparative gene expression. phyloDS is implemented as an R package and is available at https://github.com/aLiehrmann/phyloDS.

## 4 Materials and Methods

### 4.1 Phylogenetic model for the differential usage of splice junctions

#### Definition of orthologous splice junctions

Orthologous splice junctions (OSJ) are defined within the coordinate system of a multiple sequence alignment (MSA). Each MSA position (i.e., alignment column) corresponds to a set of orthologous genomic coordinates across species, except in the presence of gaps. An OSJ is defined as a pair of MSA positions corresponding to orthologous donor and acceptor splice sites, and is retained only if supported by RNA-seq evidence in at least two species. We denote by *Y_j,i,_*_in_ the number of reads supporting OSJ *j* (1 *≤ j ≤ J*) in sample *i* (1 *≤ i ≤ N*). This count is set to zero when either the donor or acceptor site is gapped in the MSA for the species from which sample *i* originates, or when the junction is not observed in that sample.

#### Simplified representation of alternative splicing across species

RNA-seq data typically reveal multiple splice junctions reflecting alternative transcripts. Rather than modeling full transcript structures, we treat individual OSJ as elementary units of alternative splicing variation. Each OSJ is considered in turn as a *focal OSJ*. For a given focal OSJ *j*, we define competing OSJ as those whose donor–acceptor interval intersects that of the focal OSJ, including OSJ sharing a single splice site. For sample *i*, we denote by *Y_j,i,_*_out_ the total number of reads supporting such competing OSJ. The relative usage of focal OSJ *j* in sample *i* is then defined as

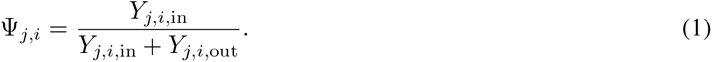

In practice, we work with its logit transformation,

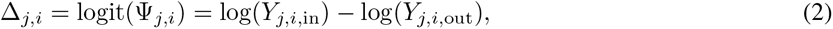

which corresponds to the log-ratio between the focal OSJ and the aggregate of competing OSJ. To ensure that the log transformation is well defined, we add a small pseudocount: *Y_j,i,k_* + *ɛ* with *ɛ* = 0.5 and *k ∈ {*in, out*}*. Our objective is to quantify how Δ*_j,i_* differs between two predefined groups of species while explicitly accounting for phylogenetic relatedness. We refer to this phylogenetically corrected group difference as the *difference-in-differences* (DID*_j_*).

#### Δ as a phylogenetic linear model

Let **Δ***_j_* = (Δ*_j,_*_1_*, . . . ,* Δ*_j,N_*)*^⊤^* denote the vector of observations for OSJ *j*. We model **Δ***_j_* using a phylogenetic linear model:

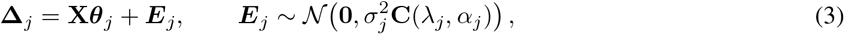

where **X** is the *N × K* design matrix, ***θ****_j_* is the vector of *K* fixed effects for OSJ *j*, and ***E****_j_* is a vector of residuals. In the two-group comparison considered throughout this work, *K* = 2. the first column of **X** is an intercept and the second is the group indicator, so that *θ_j,_*_group_ is the single coefficient of interest. The formulation is not restricted to this case. Any further covariate can be appended as an additional column of **X** in which case *K* increases accordingly. The matrix **C**(*λ_j_, α_j_*) is a correlation matrix (with unit diagonal), so that 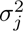 is the total variance of each component of ***E****_j_* [Ho and Ané, 2014, Gallopin et al., 2026]. The parameter *λ_j_ ∈* [0, 1] directly captures the within species correlation. The parameter *α_j_* controls the rate at which evolutionary correlations decay and is commonly interpreted through the half-life of the process, *t*_1*/*2*,j*_ = log(2)*/α_j_ ∈* (0, +*∞*) [Hansen, 1997]. It controls the level of between-species correlations. In particular, as illustrated in Figures S8–S9, when *λ_j_ →* 0 and *t*_1*/*2*,j*_ *→* 0, observations are independent within and across species. As *λ_j_ →* 1, observations from the same species become correlated. The phylogenetic tree is rescaled to unit height, so that *t*_1*/*2*,j*_ is expressed relative to the total depth of the tree: as *t*_1*/*2*,j*_ approaches 1 and beyond, correlations across species increasingly reflect the phylogenetic tree topology. The value 1 carries no special status. We use it as an upper reference point, and as the upper end of the grid explored in our empirical null simulations, because the regularized *t*_1*/*2*,j*_ estimated on both datasets analysed in this work fall below it. Testing for differential OSJ usage between groups amounts to testing the null hypothesis *H*_0_ : *θ_j,_*_group_ = 0, where *θ_j,_*_group_ (corresponding to DID*_j_*) denotes the fixed effect associated with the group indicator in the design matrix.

#### Normalization

Let the expected read count be written as E[*Y_j,i,k_*] = *q_j,i,k_ · s_i_*, where *s_i_* is a sample-specific size factor (sequencing depth) and *q_j,i,k_* reflects the underlying transcript abundance. Because Δ*_j,i_* is a within-sample log-ratio, the multiplicative factor *s_i_* cancels out, making Δ*_j,i_* intrinsically invariant to library size. This formulation also separates differential OSJ usage from differential gene expression: global changes in transcript abundance do not affect Δ*_j,i_*. However, when either *Y_j,i,_*_in_ or *Y_j,i,_*_out_ is low (e.g., nearly constitutive or absent OSJ), this invariance may break down. To mitigate this, we normalize counts and filter out low-coverage OSJ prior to testing (see Step 3 of *phyloDS workflow and implementation*).

### 4.2 Model inference and statistical test of DID*_j_* coefficients

phyloDS estimates and tests the model in Equation 3 using phyloDE [Gallopin et al., 2026]. Informally, at its core, phyloDE combines (i) phylogenetic linear regression [Ho and Ané, 2014] with (ii) empirical Bayes regularization [Ritchie et al., 2015]. The method extends the limmaCorr framework [Smyth et al., 2005] by replacing its block correlation structure with a reparameterized version of the OU covariance model in Equation 3. (i) For each OSJ *j*, the OU parameters *λ_j_* and a transformed version of *α_j_* are first estimated independently by restricted maximum likelihood using phylolm [Ho and Ané, 2014]. However, OSJ-specific parameter estimation on small phylogenies is inherently underpowered and can be unstable. Rather than testing each OSJ independently, phyloDE incorporates a regularization step to overcome this limitation: the estimated parameters are shrunk toward global values using a robust trimmed mean computed across a large number of OSJ. (ii) The resulting regularized covariance matrix is then used to decorrelate **Δ***_j_*, after which the DID*_j_* coefficients are evaluated using moderated *t*-statistics within the empirical Bayes framework of limma [Ritchie et al., 2015].

#### Positioning relative to existing methods

Conceptually, phyloDE can be positioned relative to existing approaches along two orthogonal dimensions: (*phylogenetic model*), that is, whether the phylogenetic tree topology is explicitly modeled; and (*cross-OSJ shrinkage*), that is, whether estimated parameters are regularized across OSJ–both of which are leveraged by phyloDE. Within this framework, limma and limmaCorr are regularized non-phylogenetic linear models (the latter accounting for within-species correlation), whereas phylolm and eve [Rohlfs and Nielsen, 2015] are phylogenetic approaches that estimate parameters independently, without cross-OSJ regularization (the latter based on a likelihood-ratio test).

### 4.3 phyloDS workflow and implementation

#### Step 1: Setting up a new analysis

The newExperiment function initializes a phyloDS analysis by assembling all required inputs. The primary input is a table of genomic coordinates for orthologous genes (one gene per species), including species name, chromosome, strand, start, and end positions. Additional required inputs are genome sequence file paths for each species, group assignments defining the biological or evolutionary contrast of interest (group 2 corresponds to the lineage(s) with the branch shift, and group 1 to the outgroup), and an ultrametric phylogenetic tree describing species relationships. For OSJ quantification, RNA-seq BAM files (reads aligned to their species-of-origin genome) and associated sample metadata must also be provided. The function returns a structured list containing all inputs, properly formatted for downstream analyses.

#### Step 2: Counting OSJ

The orthogroupDNA function extracts genomic sequences for each orthogroup and writes them as FASTA files. These are aligned using orthogroupMSA, which calls MAFFT in *auto* mode [Katoh, 2002], establishing the MSA coordinate system. In parallel, coverageSJ parses RNA-seq BAM files to extract splice junctions from CIGAR strings and counts supporting reads for each detected junction. Finally, findOSJ maps species-specific coordinates onto the MSA, identifies OSJ, and computes *Y_j,i,_*_in_ and *Y_j,i,_*_out_ based on the overlap structure defined in MSA coordinates. The output is a Bioconductor SummarizedExperiment object [Martin Morgan, 2017] containing MSA coordinates, species-specific genomic coordinates, and read counts for all samples.

#### Step 3: Testing differential OSJ usage

Per-sample size factors are estimated using the median-ratio method (estSizeFactors, via DESeq2 [Love et al., 2014]) and used to normalize read counts. OSJ are filtered to retain those where the (i) mean normalized counts across samples exceed a threshold (default: 1) for both spliced-in and and spliced-out components, and (ii) the focal OSJ, or at least one competing OSJ, is represented in all species. Log-ratios Δ*_j,i_* are then computed (using base-2 logarithm for interpretation purposes). The DID coefficient is tested using phyloDE by default; positive values indicate increased usage in group 2. *p*-values are adjusted for multiple testing via the Benjamini–Hochberg procedure [Benjamini and Hochberg, 1995]. Optionally, runSimu and plotSimu assess type I error control across (*λ, t*_1*/*2_) combinations via empirical null simulations.

#### Step 4: Annotation and visualization of differentially used OSJ

First, the annotateOSJ function maps OSJ from the MSA coordinate system back to the genomic coordinates of a designated reference species. Using the reference GTF annotation, donor and acceptor sites are matched to annotated exon boundaries, yielding the flanking exon identifiers (donorExon and acceptorExon). Exons fully contained within the OSJ span are additionally reported as skippedExon. Second, the predictDID function provides a complementary sequence-based analysis by predicting donor and acceptor splice site strengths across species using pre-trained OpenSpliceAI models [Chao et al., 2025]. Predicted strengths are computed independently for each species and summarized into donor and acceptor DID metrics (predDIDdonor and predDIDacceptor). Finally, the exportResults function generates IGV-ready outputs for interactive exploration. For each orthogroup, MSAs are exported as BAM files. OpenSpliceAI predictions are exported in BigWig format. Differential splicing results are written to a color-coded GTF file (significant DID *>* 0 in red, significant DID *<* 0 in blue, and non-significant OSJ in grey), where annotations correspond to the OSJ span. Projected species-specific annotations are provided as additional tracks. An IGV session file [Robinson et al., 2011] is automatically generated to load and organize all tracks with a single click.

### 4.4 Benchmarking

#### 4.4.1 Datasets and preprocessing

We analyzed two phylogenetically structured RNA-seq datasets. The first dataset consisted of brain samples from eight primate species, including three hominoids (*Homo sapiens*, *Pan troglodytes*, and *Gorilla gorilla*), three Old World monkeys (*Macaca mulatta*, *Macaca fascicularis*, and *Papio anubis*), and two New World monkeys (*Callithrix jacchus* and *Saimiri boliviensis*). All samples, except those from gorilla, were compiled by Recinos et al. [2024] and originally used to investigate cassette exon inclusion, including lineage-specific shifts in the hominoid clade. We complemented this dataset with gorilla samples from Xu et al. [2018]. The species tree was extracted from the vertebrate phylogeny available through the UCSC Genome Browser [Casper et al., 2026]. One-to-one gene orthologs (Table S7), GTF annotations, and reference genomes were retrieved from Ensembl release 113 [Harrison et al., 2024].

The second dataset comprised heart RNA-seq samples from four rodent species, including two mole-rats (*Heterocephalus glaber* and *Cryptomys damarensis*) and two outgroups (*Mus musculus* and *Cavia porcellus*). These data were compiled by Daunesse et al. [2025] to study coordinated shifts in gene expression and regulation during mole-rat evolution, including lineage-specific shifts along the branch leading to the clade comprising the two mole-rat species. At the time of our analysis, some samples generated by the authors were not publicly available in sequence repositories. We therefore replaced them with additional biological replicates from the studies compiled by the same authors [Bens et al., 2018, Faulkes et al., 2024]. The species tree was extracted from the mammalian phylogeny of Fritz et al. [2009], as curated in Hassler et al. [2022]. One-to-one orthologs were obtained from Daunesse et al. [2025] (Table S8), while GTF annotations and reference genomes were retrieved from Ensembl release 102 [Howe et al., 2021].

For both datasets, reads were aligned to the corresponding reference genomes using STAR in two-pass mode with default settings [Dobin et al., 2013]. To keep the multiple sequence alignment with MAFFT computationally tractable, we restricted the analysis to orthogroups in which all orthologous genes were at most 150 000 bp long. This retained 8 867 of 11 298 orthogroups in the *hominoids* dataset and 12 841 of 14 404 in the *mole-rats* dataset. Metadata for all samples (including SRA identifiers), species information (including FTP links to reference genome and GTF files), species trees, and the final lists of orthologous genes analyzed in this study are provided in the Supplementary Data (see Data availability).

#### 4.4.2 Simulation framework and comparison with reference-based methods

##### Empirical null simulations with compcodeR

To assess type I error control under the global null hypothesis, we simulated OSJ count profiles using compcodeR [Soneson, 2014] within the phylogenetic Poisson log-normal framework of Bastide et al. [2023]. Simulations were calibrated separately on the *hominoids* and *mole-rats* datasets to reproduce empirical mean–dispersion relationships estimated with DESeq2. For each OSJ, *Y_j,i,_*_in_ and *Y_j,i,_*_out_ were simulated independently under OU processes on latent log-counts, followed by Poisson sampling. The group effect was set to zero for spliced-in and spliced-out components, ensuring DID*_j_* = 0 for all simulated OSJ. We explored a grid of parameters: *λ ∈ {*0, 0.1, 0.25, 0.5, 0.75, 0.9*}* (from independent replicates to species-structured) and *t*_1*/*2_ *∈ {*0.05, 0.1, 0.2, 0.5, 1*}* (from weak to strong phylogenetic constraint), where *t*_1*/*2_ is expressed relative to unit tree height. For each parameter combination, we simulated 5 000 OSJ profiles and repeated the procedure 50 times. For each replicate, the empirical type I error rate was computed as 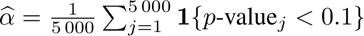. Under a well-calibrated procedure, *α̂* is expected to be close to *α*; we report averages across replicates. This simulation procedure is implemented in phyloDS via the runSimu and plotSimu functions, allowing users to assess calibration on their own datasets.

##### Comparison with reference-based OSJ inference

We inferred OSJs using a reference-based strategy [Recinos et al., 2024]. For each dataset, all RNA-seq reads were aligned to the genome of the reference species using STAR in two-pass mode: the *Homo sapiens* genome for the *hominoids* dataset and the *Mus musculus* genome for the *mole-rats* dataset. OSJs were then defined in the coordinate system of the reference genome and retained when supported by at least two species.

##### Comparison with reference-based differential splicing event detection

We generated annotated splicing events from the reference species of each dataset (*Homo sapiens* for *hominoids*, *Mus musculus* for *mole-rats*) using SUPPA2 [Trincado et al., 2018] together with the corresponding GTF annotation. Throughout, we follow the SUPPA2 event nomenclature: skipping exon (SE), alternative 5’ and 3’ splice sites (A5, A3), mutually exclusive exons (MX), and alternative first and last exons (AF, AL). The resulting events were then projected onto the MSA coordinate system to define orthologous events across species. For each event, we constructed a count matrix from the underlying OSJ counts following event-type-specific aggregation rules. For events involving two junctions only (A3, A5, AF, AL), one junction was arbitrarily assigned to the spliced-in component and the other to the spliced-out component. For SE events, the spliced-in component was defined as the average of the two junctions flanking the skipped exon, and the spliced-out component as the exon-skipping junction. For MX events, the spliced-in and spliced-out components were defined as the average of the two pairs of junctions flanking the first and second mutually exclusive exons, respectively. Differential usage was then tested with the same procedure as in phyloDS via phyloDE. To enable comparison between the two strategies, we counted significant events for phyloDS as follows: an annotated event was considered significant if at least one of its constituent OSJ was significant (*p*_adj_ *<* 0.1); the number of significant unannotated events was estimated as the number of mutually non-overlapping significant OSJ that did not overlap any annotated event.

#### 4.4.3 Validation of significant OSJ

##### Comparison with OpenSpliceAI–inferred 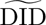

We extracted the genomic sequences of each orthologous gene and, using OpenSpliceAI [Chao et al., 2025], predicted splice site strengths with a pre-trained reference model (trained on *Homo sapiens* for the *hominoids* dataset and on *Mus musculus* for the *mole-rats* dataset). For each species *i* and OSJ *j*, OpenSpliceAI returns predicted splice site strength scores *p_j,i,_*_acceptor_ and *p_j,i,_*_donor_, corresponding to the acceptor and donor sites, respectively. To quantify sequence-based differences between the two groups defined within each dataset, we computed a predicted difference-in-differences statistic (denoted 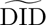_*j*,acceptor_ and 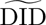_*j*,donor_) as the difference in logit-transformed mean splice site strengths between groups:

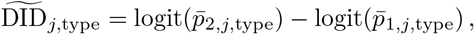

where *p̅*_1,*j*,type_ and *p̅*_2,*j*,type_ denote the mean predicted splice site strengths across species in groups 1 and 2, respectively, and type *∈ {*acceptor, donor*}*. In practice, the logit transform was computed using base-2 logarithms, i.e. logit(*x*) = log_2_(*x/*(1 *− x*)). Finally, for each OSJ, we retained the effect with the largest absolute magnitude:

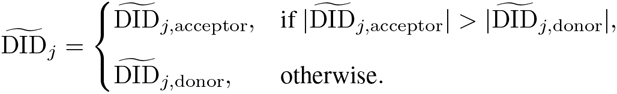

Positive 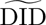_*j*_ indicates stronger predicted splice site strength in group 2 relative to group 1, whereas negative values indicate stronger predicted splice sites in group 1. Retaining the larger-magnitude site rather than averaging the two is supported by the joint distribution of donor and acceptor effects (Figure S12), which is concentrated along the two axes: when one site shows a strong predicted shift, the other typically remains near zero, so averaging would dilute the signal. Furthermore, to avoid pseudo-replication, we further grouped OSJs into loci by merging overlapping intron spans and retained, per locus, the single OSJ with the largest *|*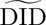_*j*_*|* as its representative.

The procedure to compute 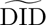*_j_*,_acceptor_ and 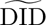*_j_*,_donor_ is implemented in phyloDS via the predictDID functions, allowing users to generate sequence-based predictions directly on their own datasets using the available models for *Homo sapiens*, *Mus musculus*, *Danio rerio*, and *Arabidopsis thaliana*. Although OpenSpliceAI is not inherently mechanistic, it can provide independent sequence-level support for the observed shifts.

#### Functional enrichment analysis

We extracted Ensembl gene identifiers from the reference species (*Homo sapiens* for *hominoids* and *Mus musculus* for *mole-rats*) for genes harboring at least one OSJ identified by phyloDS. Functional enrichment analyses were subsequently performed separately for the *hominoids* and *mole-rats* datasets using gprofiler2 [Kolberg et al., 2020]. Enrichment was tested across the annotation sources provided by gprofiler2 using the built-in FDR correction (Benjamini–Hochberg). Terms were considered significant at *p*_adj_ *<* 0.05. For visualization in Figure 2.C, we retained a subset of representative, non-redundant terms and pathways summarizing the main functional signals observed in each dataset.

#### 4.4.4 Runtime and memory usage of phyloDS

The complete analysis workflow is orchestrated using a dedicated Nextflow pipeline (Di Tommaso et al. [2017]; see Data Availability), which integrates both pre-processing steps (data download, read alignment) and downstream analyses using the phyloDS R package. This implementation ensures efficient execution on high-performance computing (HPC) infrastructures. All computations were performed on *AMD EPYC Genoa* and *AMD EPYC Milan* processors.

Constructing the MSA is by far the dominant cost, accounting for more than 97% of the total phyloDS runtime in both datasets (*∼*189 CPU hours for *hominoids* and *∼*450 CPU hours for *mole-rats*). This cost is, however, embarrassingly parallel: each of the 8 867 *hominoids* and 12 841 *mole-rats* orthogroups is aligned independently in a short, low-memory task (median *∼*20 s, using less than 1 GB of memory for the vast majority of orthogroups; the longest loci reach *∼*45 min and stay below 9 GB). Distributed across 30 concurrent alignment jobs, the full set of alignments is therefore produced in roughly 6 (*hominoids*) to 15 (*mole-rats*) hours of wall-clock time. Crucially, the MSAs depend only on the set of species and their genome assemblies, not on the RNA-seq data: they are a one-time investment that can be reused across any number of subsequent datasets sharing the same assemblies.

Once the MSAs of orthologous genes are available, the comparative analysis itself is not the bottleneck. Counting splice junctions from the aligned BAM files is the memory-limiting step (peak *∼*50 GB for *mole-rats* and *∼*40 GB for *hominoids*), yet it runs as a single task in about one hour of wall-clock time. All subsequent steps complete in minutes to about an hour on a single node and never exceed 30 GB. End to end, the downstream statistical analysis of a full dataset therefore requires only a few hours of wall-clock time (*∼*3 h).

## 5 Data Availability

### Software

The phyloDS method is implemented as an R package and is publicly available at https://github.com/aLiehrmann/phyloDS. The repository provides installation instructions, documentation, and a detailed vignette.

### Data and analyses

All data used in the *hominoids* and *mole-rats* analyses were obtained from publicly available sources, mapped, and then analyzed using phyloDS. The complete analysis workflow (including automated data retrieval) is implemented in a dedicated Nextflow pipeline, available at https://github.com/aLiehrmann/phyloDS_reproducibility. All figures can be reproduced using the scripts provided in the same repository.

## 6 Supplementary Material

Supplementary Material contains exploratory quality-control analyses of both datasets (Figure S1); IGV sessions illustrating differentially used OSJs that involve unannotated splice sites in representative genes (Figures S2–S7); tree transformations showing how the Ornstein–Uhlenbeck parameters *λ* and *t*_1*/*2_ shape the phylogenetic correlation structure (Figures S8–S9); the type I error rate of phylolm under the global null (Figure S10); sequence-based control analyses for non-differentially used OSJs and the joint distribution of donor and acceptor predicted effects (Figures S11–S12); and the complete lists of orthologous genes, differentially used OSJs, unannotated OSJs, and functional enrichment results for both datasets (Tables S1–S8).

## 7 Acknowledgments

This work was granted access to the HPC resources of the MeSU platform at Sorbonne Université. The MeSU platform and its supercomputing resources are supported by the Ile-de-France region, the French Ministère de l’Enseignement Supérieur et de la Recherche, and Sorbonne Université. This work has been funded by the European Union (ERC, PROMISE, 101087830). Views and opinions expressed are however those of the author(s) only and do not necessarily reflect those of the European Union or the European Research Council. Neither the European Union nor the granting authority can be held responsible for them. For the purpose of open access, a CC-BY public copyright licence has been applied by the authors to the present document and will be applied to all subsequent versions up to the Author Accepted Manuscript arising from this submission. This work has also been co-funded by the French National Agency for Research (ANR, France 2030, PostGenAI@Paris, ANR-23-IACL-0007).

## A Exploratory analyses

**Figure S1:**
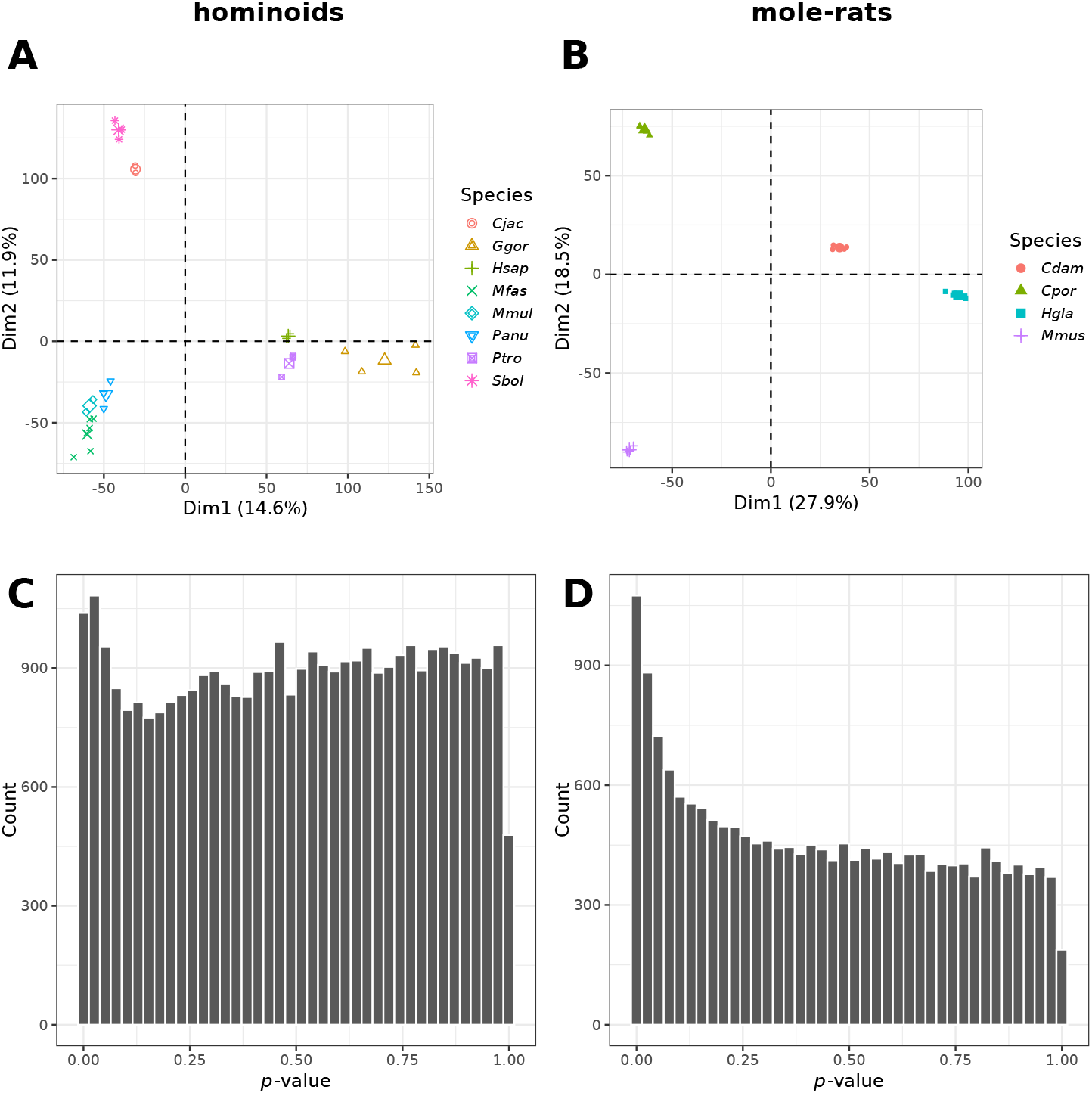
Exploratory analyses of the *hominoids* and *mole-rats* datasets. (**A**,**B**) Principal component analysis (PCA) of orthologous OSJ-level log-ratio values Δ*_j,i_* after normalization of the counts (see the *Phylogenetic model for the differential usage of splice junctions* section in Methods). Each point corresponds to a biological replicate (larger symbols indicate species means). Projection onto the first principal component, explaining 14.6% and 27.9% of the variance for *hominoids* and *mole-rats*, respectively, reveals a clear separation between the predefined clades. In *hominoids*, group 1 (*Macaca fascicularis*, *Mfas*; *Macaca mulatta*, *Mmul*; *Papio anubis*, *Panu*; *Callithrix jacchus*, *Cjac*; *Saimiri boliviensis*, *Sbol*) is separated from group 2 (*Gorilla gorilla*, *Ggor*; *Homo sapiens*, *Hsap*; *Pan troglodytes*, *Ptro*). In *mole-rats*, group 2 (*Cryptomys damarensis*, *Cdam*; *Heterocephalus glaber*, *Hgla*) is distinct from the outgroup species (*Cavia porcellus*, *Cpor*; *Mus musculus*, *Mmus*), consistent with the study design. (**C**,**D**) Histograms of raw *p*-values from the differential splicing analysis. Both datasets show enrichment of small *p*-values, indicative of true signal, while the bulk of the distribution remains approximately uniform, with some deviations in the extreme right tail. Overall, these exploratory analyses do not reveal clear anomalies that would compromise downstream inference.

## B IGV sessions of differentially used OSJs involving unannotated splice sites

**Figure S2:**
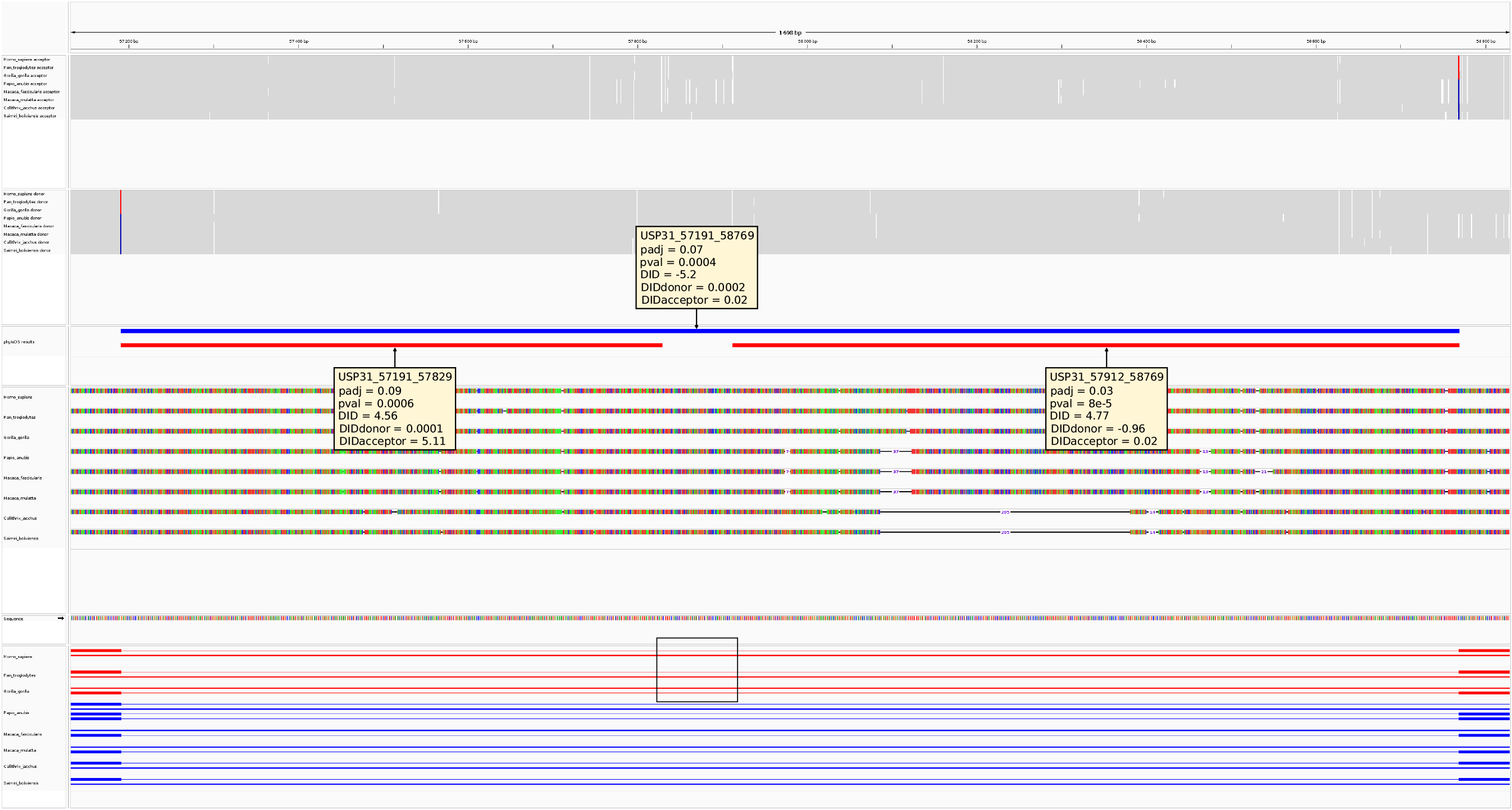
IGV session exported by phyloDS illustrating an unannotated skipping exon (SE) event in *USP31* detected in the *hominoids* dataset (see Figure 3 for guidance on how to interpret the tracks). The exon is not annotated in any of the species in the hominoids group 2 (see dark rectangle), in which it is relatively more used than in non-hominoid primates (group 1), as indicated by phyloDS (DID *>* 4.5; *p*_adj_ *<* 0.1) and OpenSpliceAI (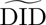_acceptor_ > 5, retaining the largest effect size in the region). Note that this strong 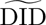 is not incompatible with the faint donor/acceptor strength tracks, since 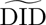 is a difference of logits.

**Figure S3:**
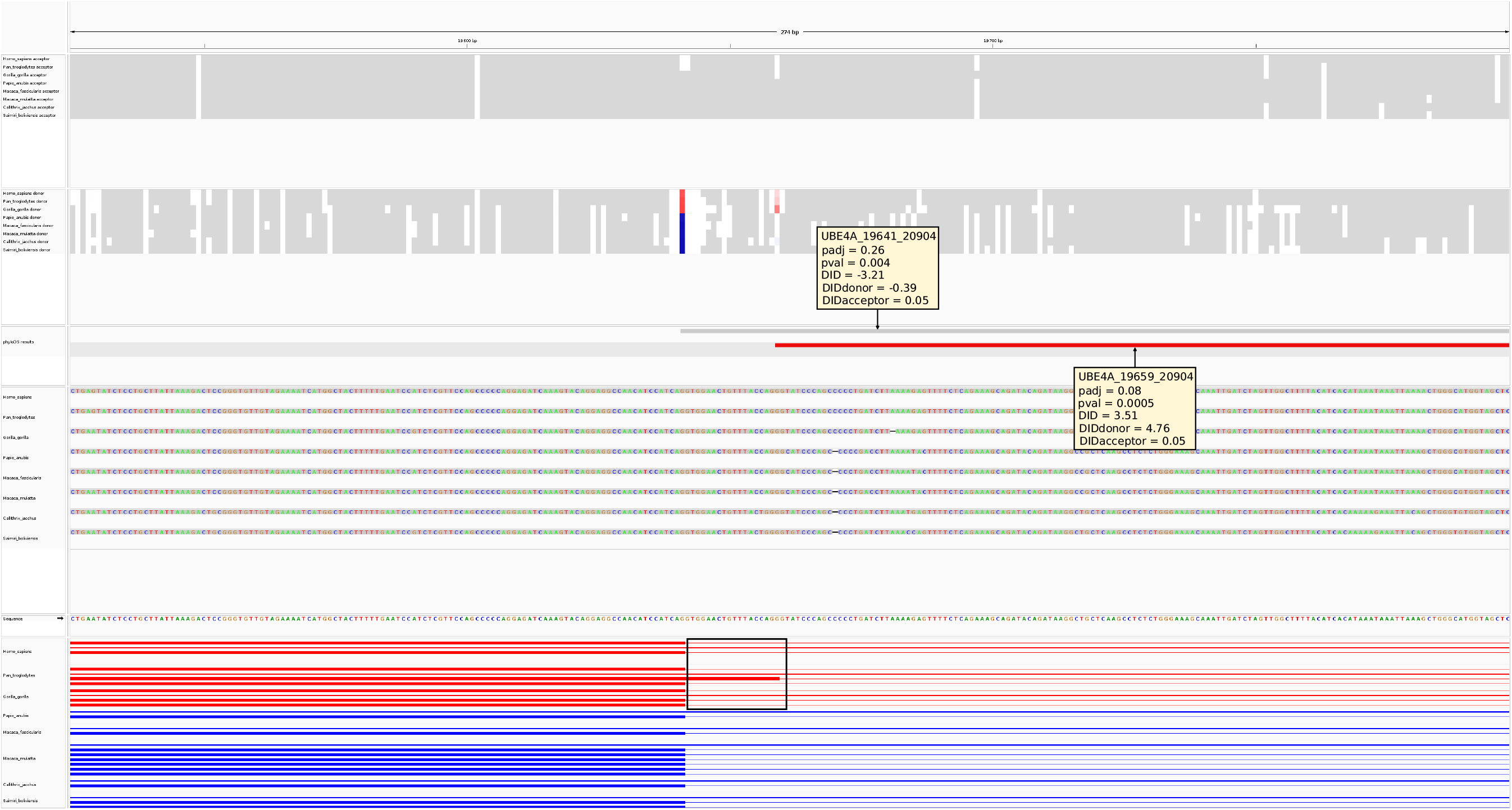
IGV session exported by phyloDS illustrating an unannotated alternative 5’ splice site (A5) event in *UBE4A* detected in the *hominoids* dataset (see Figure 3 for guidance on how to interpret the tracks). The second 5’ splice site is annotated only in *Pan troglodytes* within the hominoids group 2 (see dark rectangle), where it is relatively more used than in non-hominoid primates (group 1), as indicated by phyloDS (DID = 3.51; *p*_adj_ = 0.08) and OpenSpliceAI (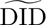_donor_ = 4.76, retaining the largest effect size in the region).

**Figure S4:**
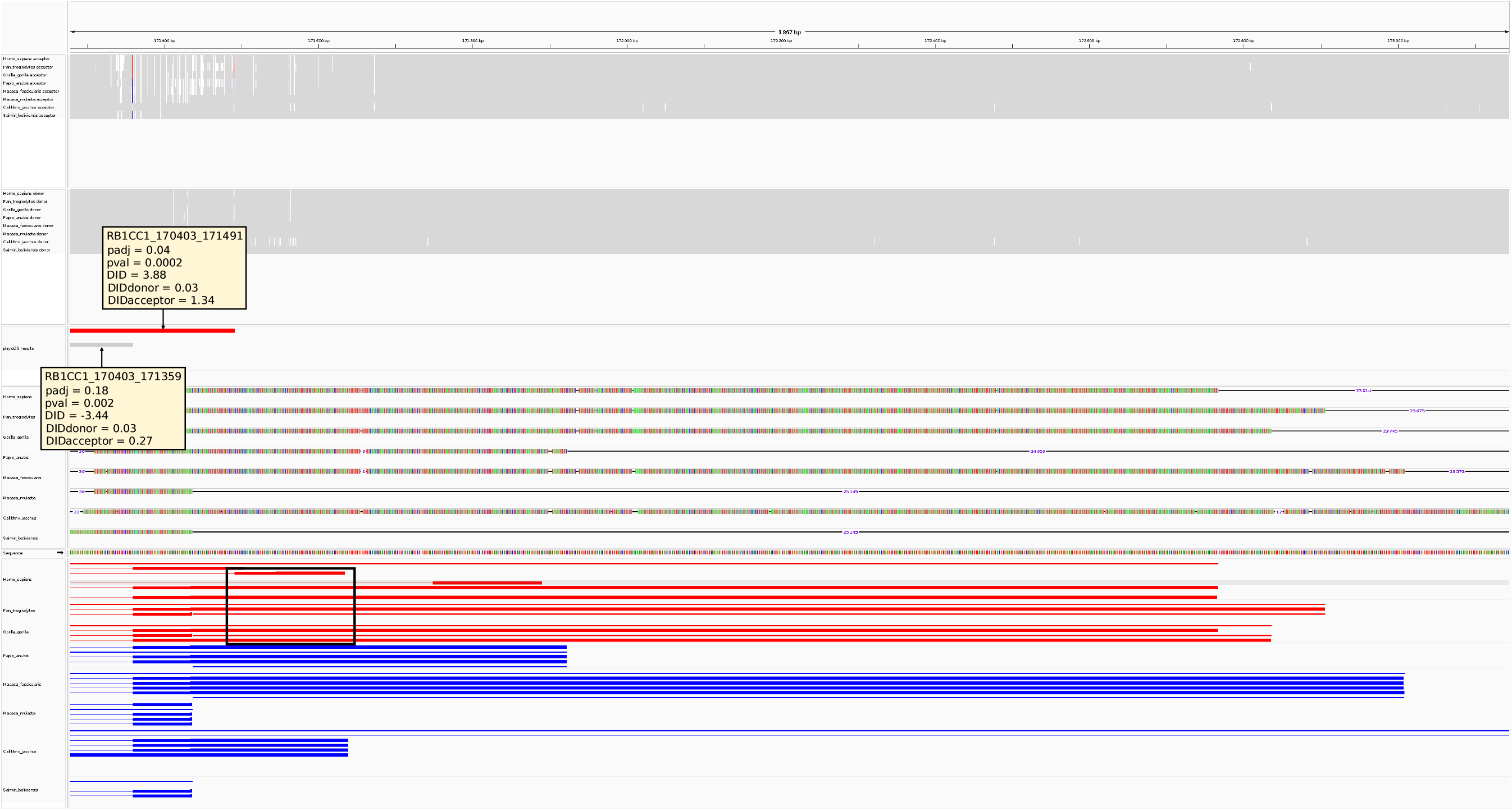
IGV session exported by phyloDS illustrating an unannotated alternative last exon (AL) event in *RB1CC1* detected in the *hominoids* dataset (see Figure 3 for guidance on how to interpret the tracks). A last exon is annotated only in *Homo sapiens* within hominoids group 2 (see dark rectangle), where it is relatively more used than in non-hominoid primates (group 1), as indicated by phyloDS (DID = 3.88; *p*_adj_ = 0.04) and OpenSpliceAI (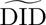_acceptor_ = 1.34, retaining the largest effect size in the region).

**Figure S5:**
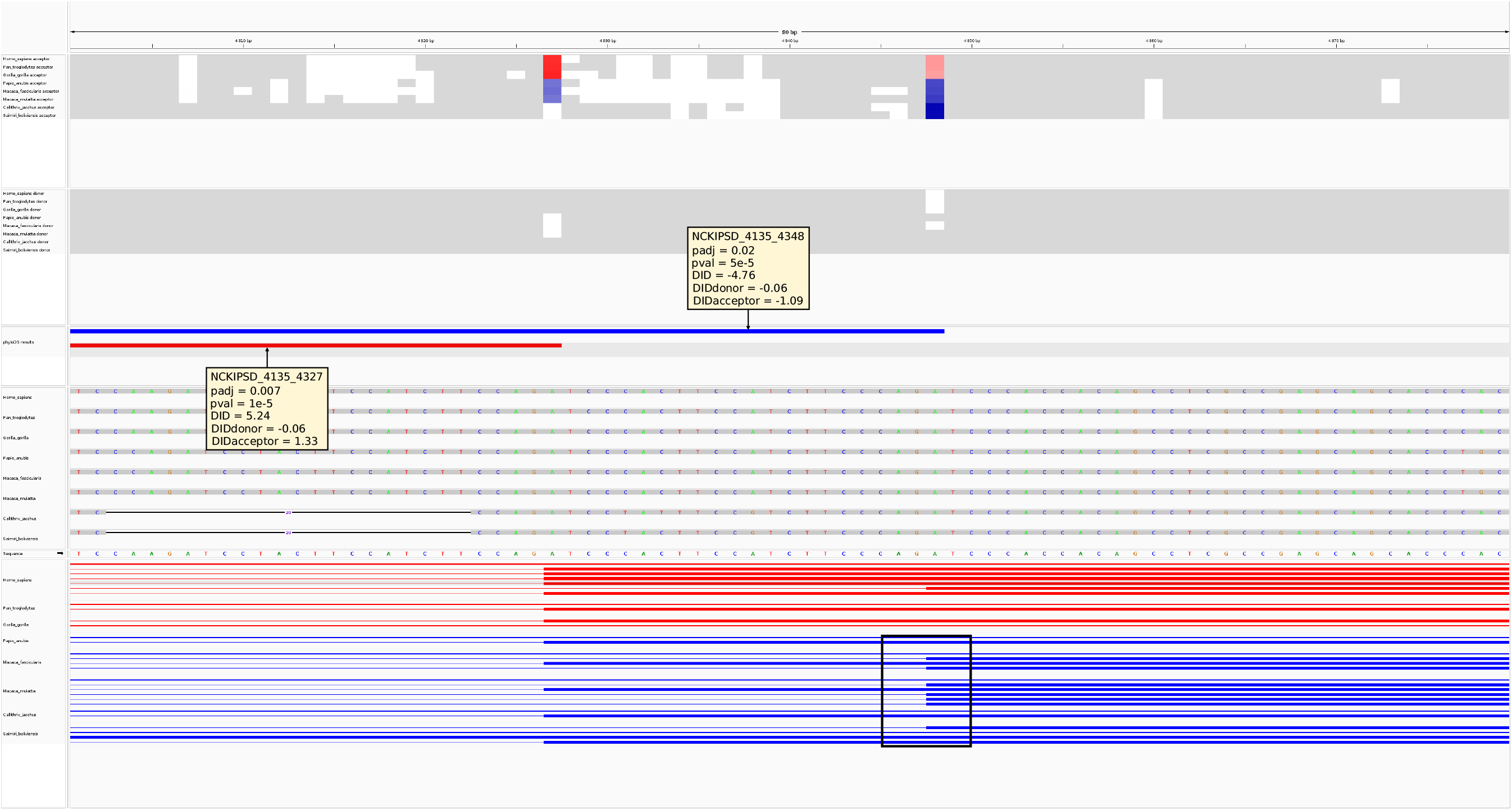
IGV session exported by phyloDS illustrating an unannotated alternative 3’ splice site (A3) event in *NCKIPSD* detected in the *hominoids* dataset (see Figure 3 for guidance on how to interpret the tracks). The second 3’ splice site is not annotated in *Papio anubis* and *Callithrix jacchus* within the non-hominoid primates group 1 (see dark rectangle), where it is relatively more used than in hominoids (group 2), as indicated by phyloDS (DID = *−*4.76; *p*_adj_ = 0.02) and OpenSpliceAI (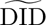_acceptor_ = 1.09, retaining the largest effect size in the region).

**Figure S6:**
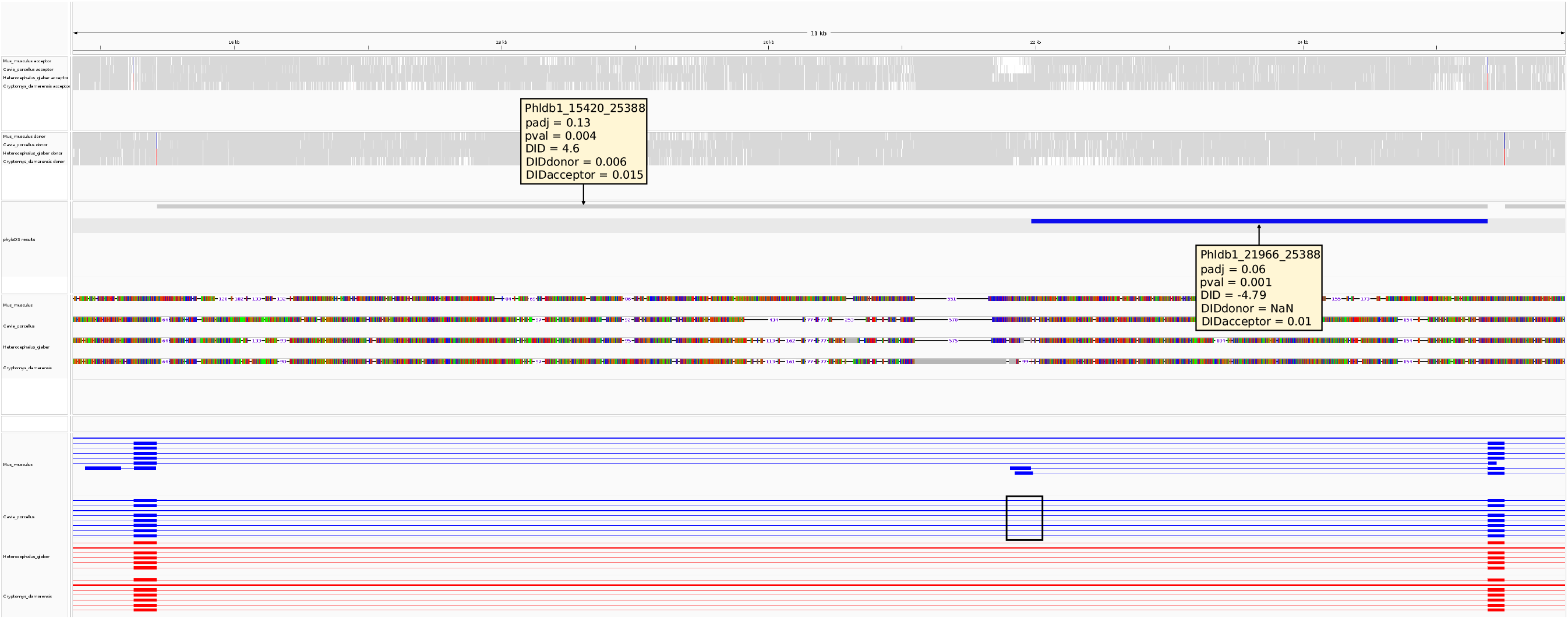
IGV session exported by phyloDS illustrating an unannotated alternative first (AF) exon event in *Phldb1* detected in the *mole-rats* dataset (see Figure 3 for guidance on how to interpret the tracks). A first exon is annotated in *Mus musculus* and not in *Cavia porcellus* (see dark rectangle), both belonging to the outgroup 1, where it is relatively more used than in mole-rats (group 2), as indicated by phyloDS (DID = *−*4.76; *p*_adj_ = 0.06).

**Figure S7:**
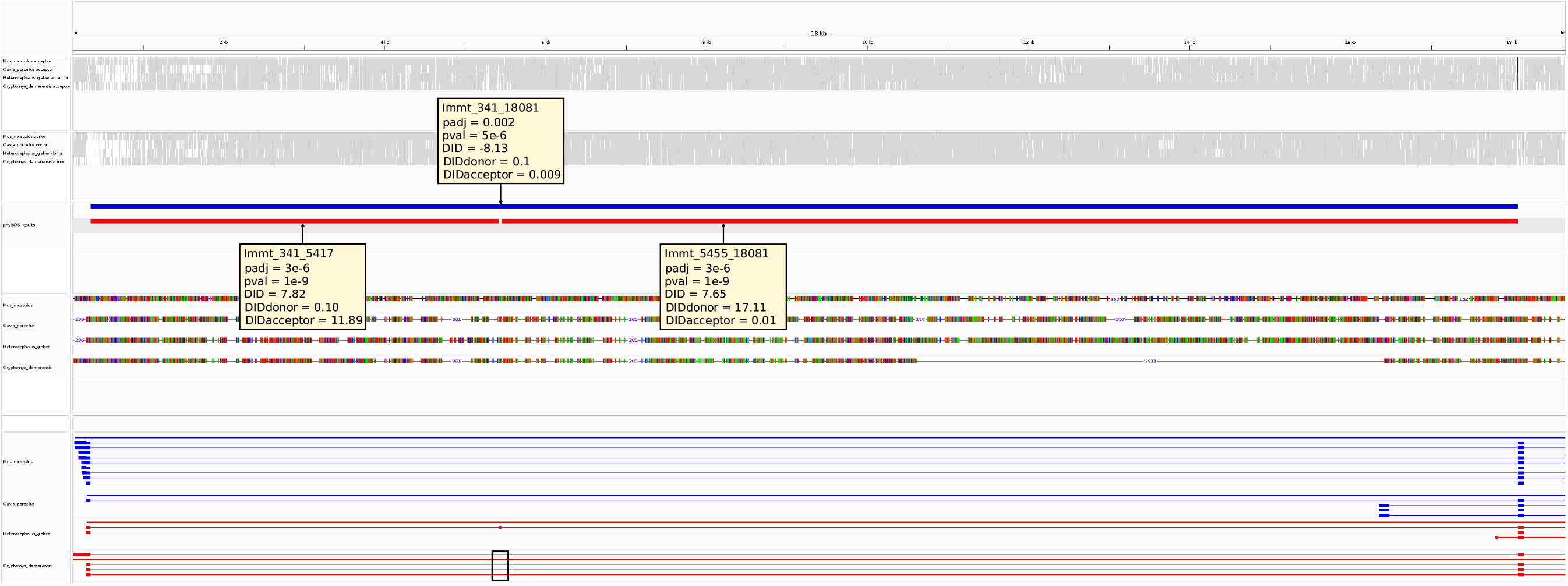
IGV session exported by phyloDS illustrating an unannotated skipping exon (SE) event in *Immt* detected in the *mole-rats* dataset (see Figure 3 for guidance on how to interpret the tracks). The exon is annotated in *Heterocephalus glaber* and not in *Cryptomys damarensis* (see dark rectangle), both belonging to the mole-rats group 2, in which it is relatively more used than other rodents (group 1), as indicated by phyloDS (DID *>* 7.5; *p*_adj_ *<* 10*^−^*^5^) and OpenSpliceAI (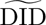_acceptor_ and 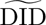_donor_ > 10, retaining the largest effect size in the region).

### C Tree transformations under the OU model

**Figure S8:**
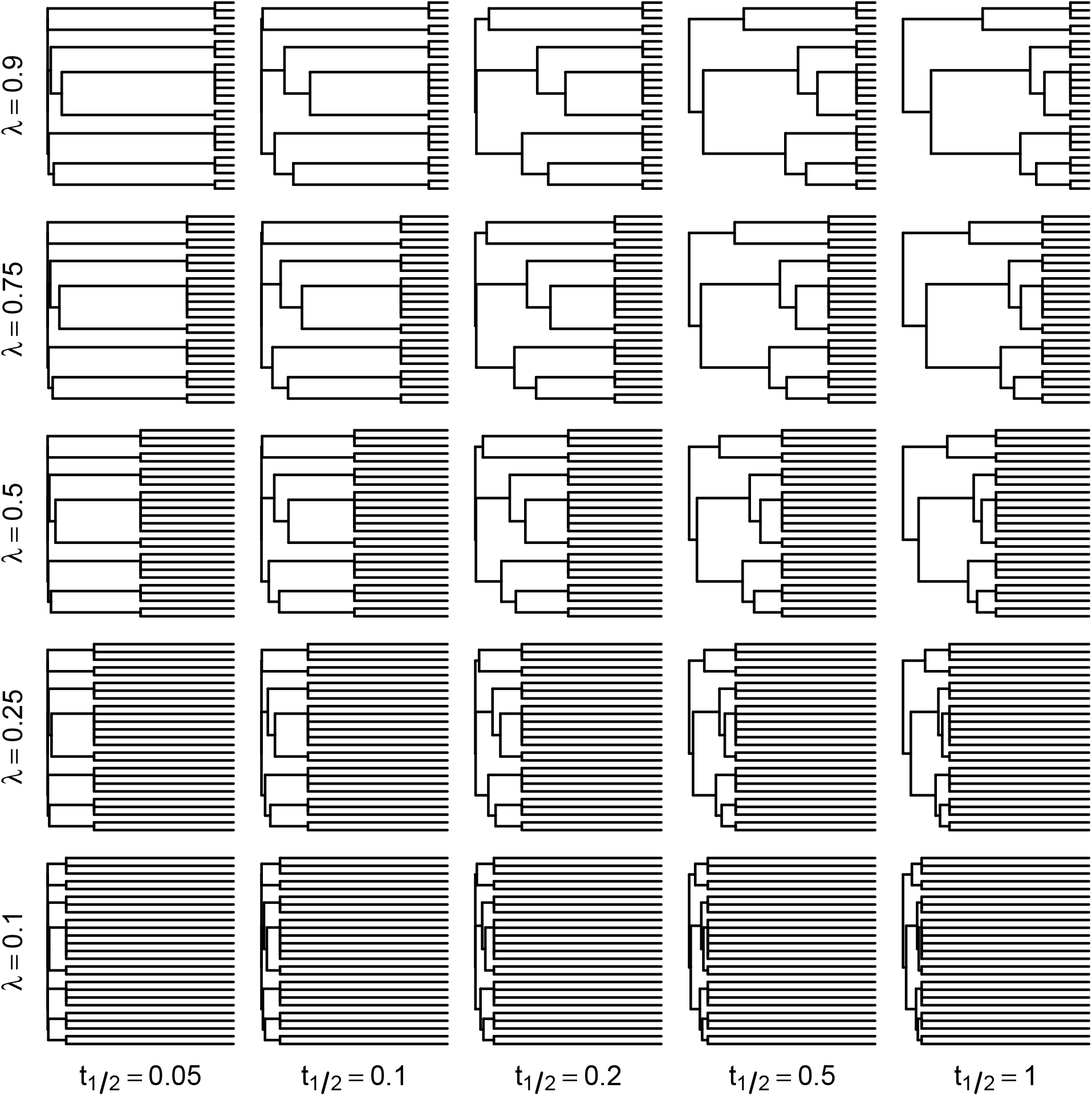
Tree transformations illustrating how *λ* and *t*_1*/*2_ shape the correlation structure in the *hominoids* dataset. A Brownian motion (BM) evolving on each transformed tree induces the same covariance matrix as the Ornstein–Uhlenbeck model of Equation 3. When *λ* decreases, biological replicates become more independent. When *t*_1*/*2_ decreases, the tree tends toward a star tree.

**Figure S9:**
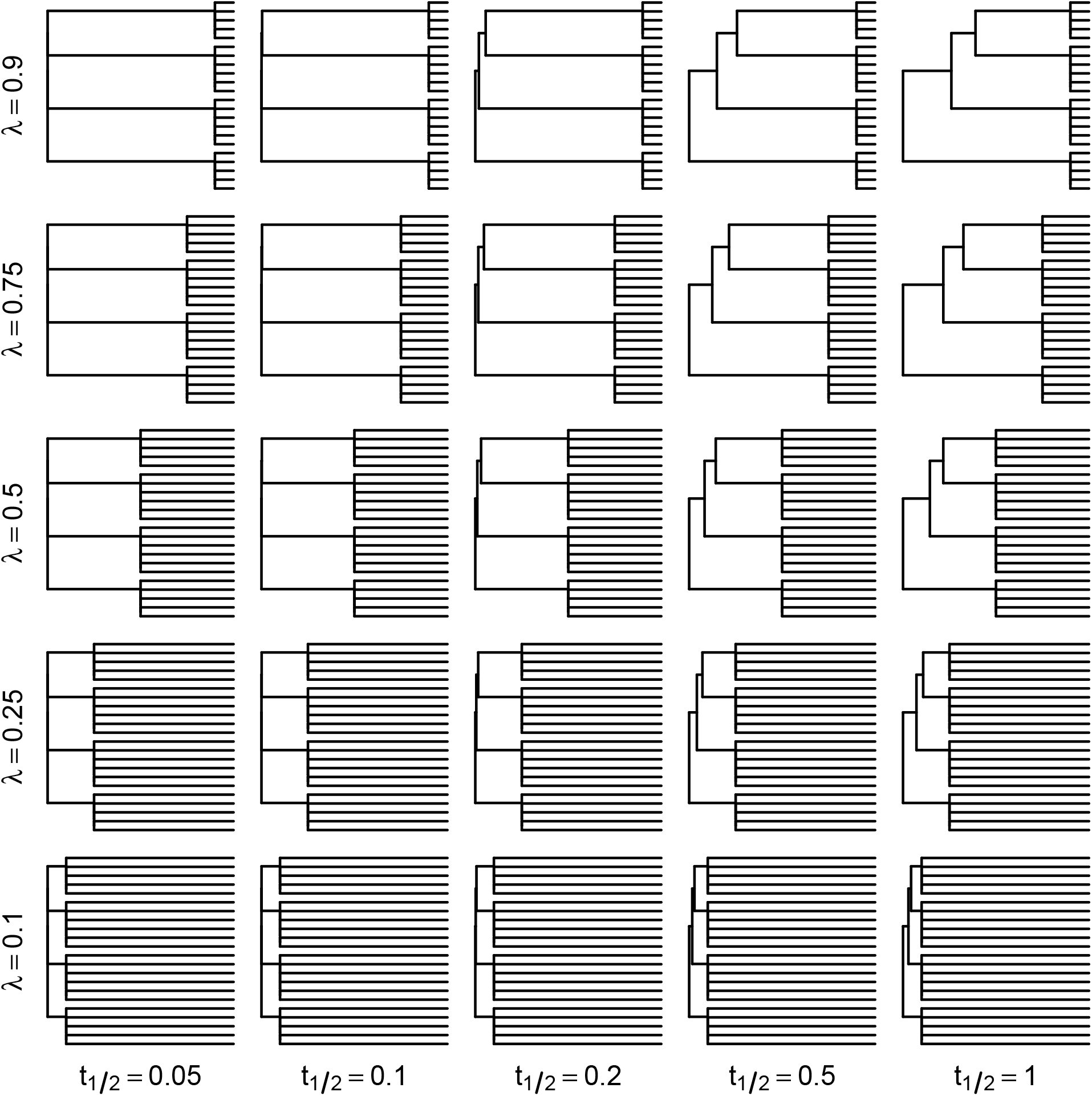
Tree transformations illustrating how *λ* and *t*_1*/*2_ shape the correlation structure in the *mole-rats* dataset. A Brownian motion (BM) evolving on each transformed tree induces the same covariance matrix as the Ornstein–Uhlenbeck model of Equation 3. When *λ* decreases, biological replicates become more independent. When *t*_1*/*2_ decreases, the tree tends toward a star tree.

### D Type I error rate of limmaCorr and phylolm

**Figure S10:**
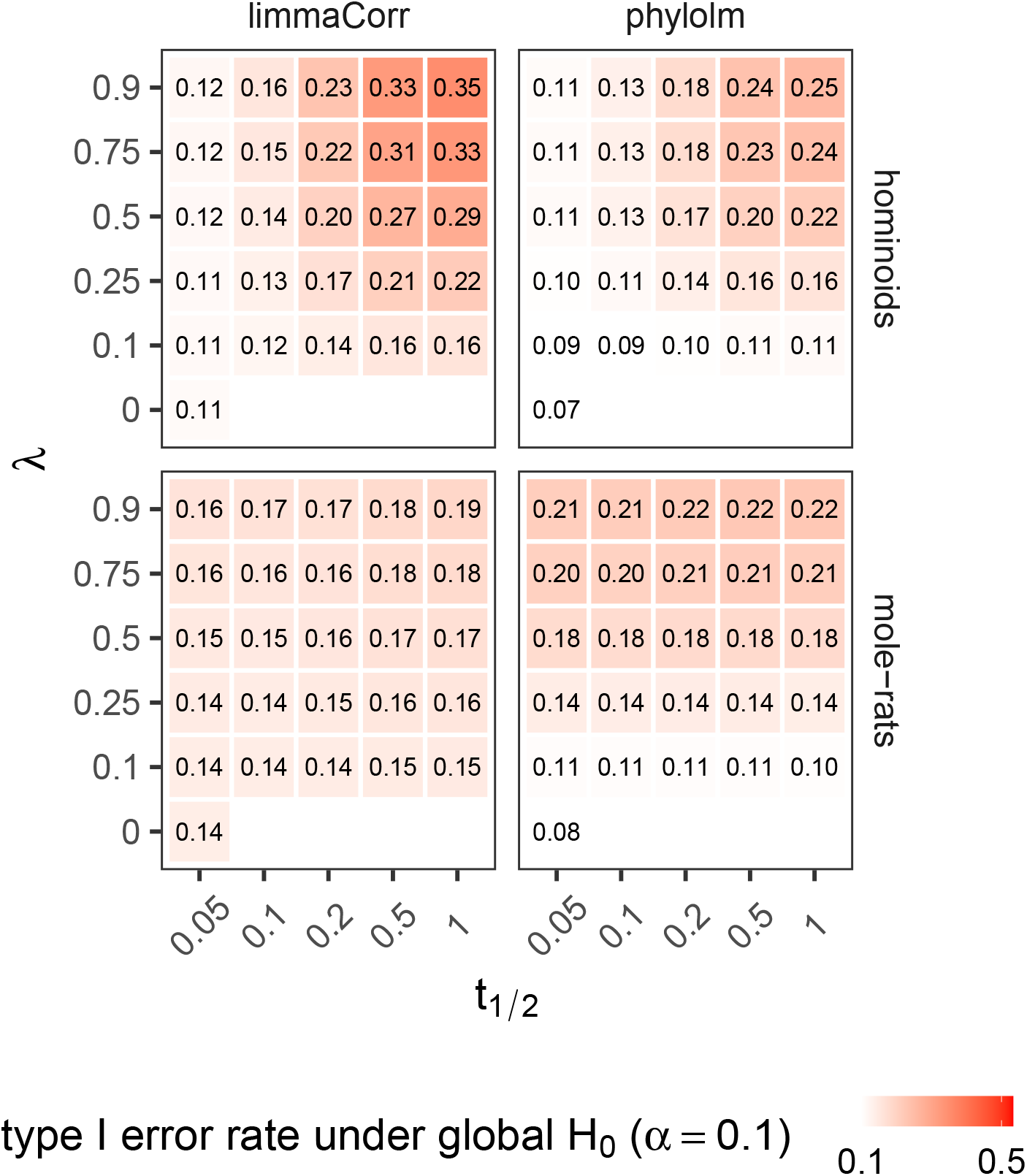
Type I error rate under the global null hypothesis (*H*_0_) at nominal level *α* = 0.1 for limmaCorr and phylolm. limmaCorr is a regularized non-phylogenetic linear model approach accounting for within-species variation, and phylolm is a phylogenetic approach that estimates parameters independently for each OSJ. OSJ counts (spliced-in and spliced-out) were simulated with compcodeR under a phylogenetic Poisson log-normal model calibrated on the *hominoids* and *mole-rats* datasets to match empirical mean–dispersion relationships. For each scenario, 5 000 OSJs were simulated with no group effect (DID = 0), across a grid of phylogenetic signals (*λ*; ranging from independent replicates to species-structured variation) and OU half-lives (*t*_1*/*2_; ranging from independent species to tree-structured variation), with 50 replicates. The empirical type I error rate was computed as the proportion of raw *p*-values below *α* and averaged across replicates.

### E Sequence-based support with OpenSpliceAI

**Figure S11:**
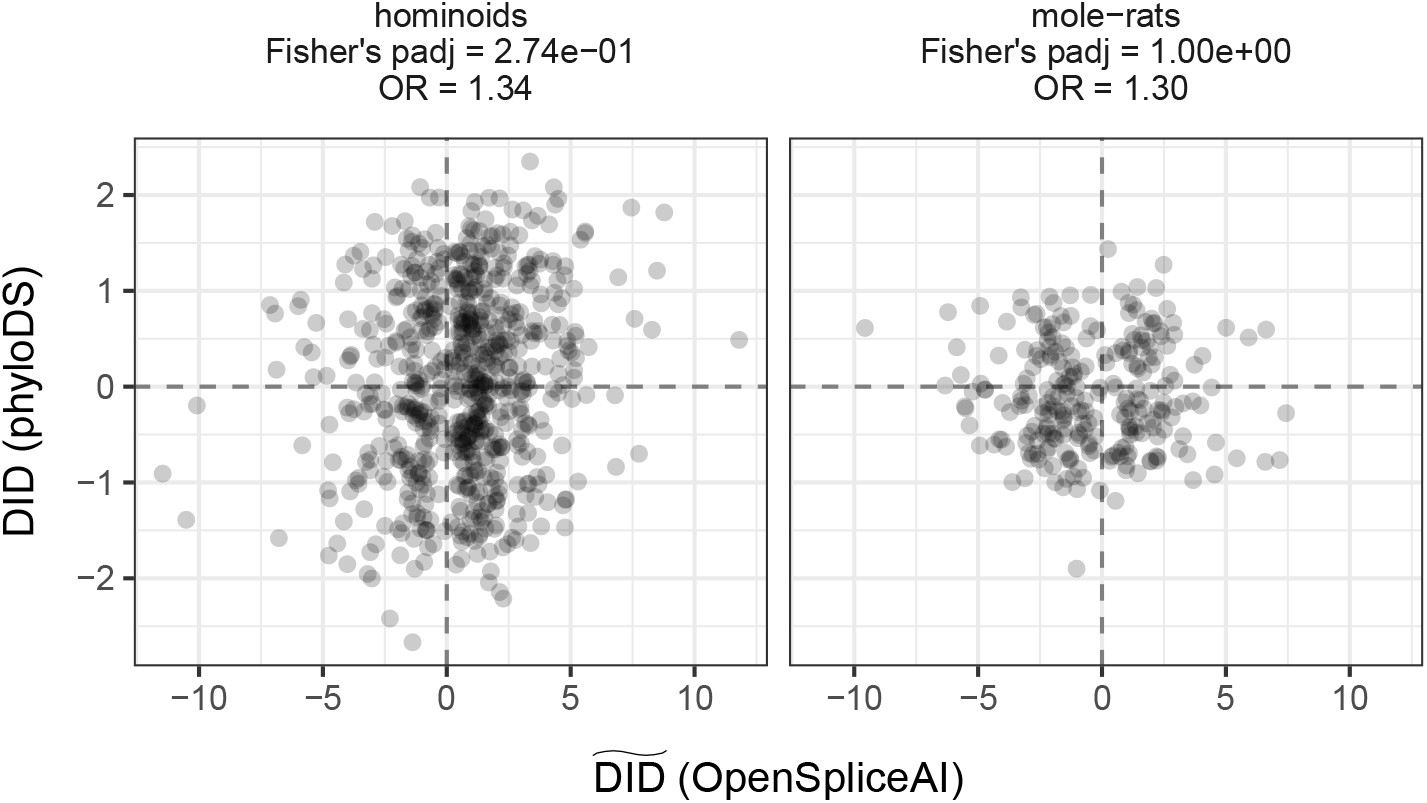
Comparison of difference-in-differences estimates derived from transcriptomic data using phyloDS and from sequence-based predictions using OpenSpliceAI. Each point represents an OSJ, with DID estimates plotted against OpenSpliceAI-predicted 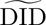 scores. Panels are stratified by dataset (*hominoids* and *mole-rats*). Only clearly non-significant OSJs according to phyloDS (*p*_padj_ *>* 0.9) are shown here. For these OSJs, concordance between transcriptomic and sequence-based effect directions is not significant in either dataset according to Fisher’s exact test (*p*_padj_ and odds ratios are reported in each panel).

**Figure S12:**
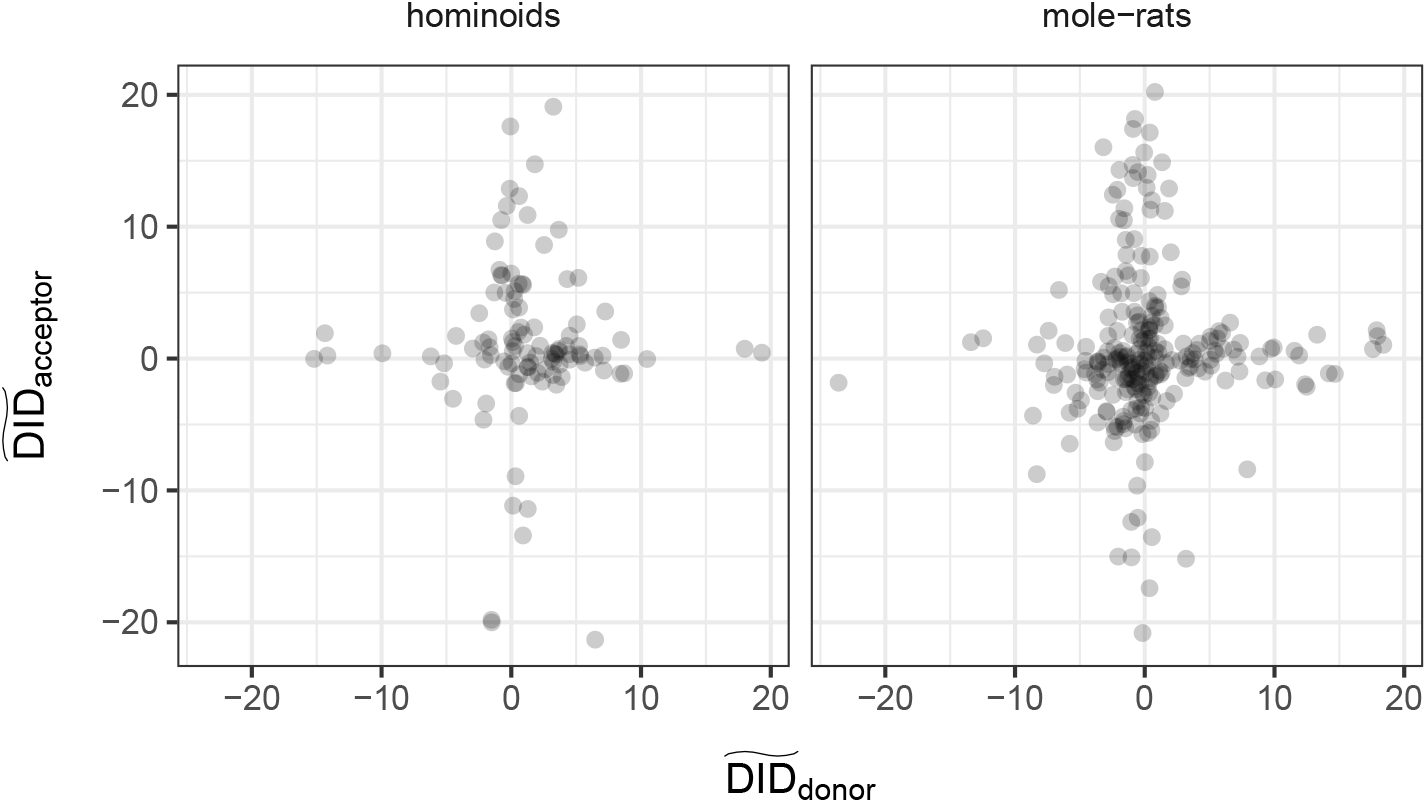
Donor and acceptor sequence-based effects are rarely strong simultaneously. For a set of representative differentially used OSJs, the predicted difference-in-differences at the donor site (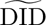*_j,_*_donor_, *x*-axis) is plotted against that at the acceptor site (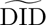*_j,_*_acceptor_, *y*-axis). Panels are stratified by dataset (*hominoids* and *mole-rats*). Both distributions are concentrated along the two axes rather than the diagonals: when one splice site shows a strong predicted shift, the other typically remains near zero. This pattern motivates summarizing each OSJ by the site with the larger absolute effect, as averaging the two would dilute a signal carried by a single site.

